# Genome-wide identification and analysis of prognostic features in human cancers

**DOI:** 10.1101/2021.06.01.446243

**Authors:** Joan C. Smith, Jason M. Sheltzer

## Abstract

Clinical decisions in cancer rely on precisely assessing patient risk. To improve our ability to accurately identify the most aggressive malignancies, we constructed genome-wide survival models using gene expression, copy number, methylation, and mutation data from 10,884 patients with known clinical outcomes. We identified more than 100,000 significant prognostic biomarkers and demonstrate that these genomic features can predict patient outcomes in clinically-ambiguous situations. While adverse biomarkers are commonly believed to represent cancer driver genes and promising therapeutic targets, we show that cancer features associated with shorter survival times are not enriched for either oncogenes or for successful drug targets. Instead, the strongest adverse biomarkers represent widely-expressed housekeeping genes with roles in cell cycle progression, and, correspondingly, nearly all therapies directed against these features have failed in clinical trials. In total, our analysis establishes a rich resource for prognostic biomarker analysis and clarifies the use of patient survival data in preclinical cancer research and therapeutic development.

## Introduction

The ability to accurately discriminate between aggressive and indolent cancers underlies the prediction of patient risk and can guide crucial treatment decisions^1^. For benign cancers, watchful waiting and/or surgical resection can be appropriate, while invasive cancers may require multimodal treatment with cytotoxic therapies that themselves cause substantial morbidity. Both cancer under-treatment and cancer-overtreatment have been identified as significant sources of patient mortality, underscoring the urgent need to improve our ability to precisely identify patients with the most aggressive malignancies^2–7^.

Current risk prediction largely relies on histopathological and radiological assessment of disease status^1^. The presence of features like lymph node metastases and cellular dedifferentiation have been identified as strong predictors of patient outcome and are used to determine cancer stage and grade^8^. However, these pathological markers require subjective judgements and can exhibit low levels of interobserver agreement^9–14^. Moreover, even perfect tumor staging cannot unambiguously predict a patient’s subsequent clinical course^15–17^.

The widespread adoption of gene expression analysis, DNA sequencing, copy number determination, and other genomic technologies in clinical settings has raised the exciting possibility that molecular markers could be developed to improve risk assessment^18^. For instance, a 21-gene RT-PCR panel called Oncotype DX has been demonstrated to accurately predict the likelihood of disease recurrence in ER+ breast cancer, and assigning chemotherapy to patients identified as high-risk based on Oncotype DX decreases the frequency of disease recurrence^19^. Similar gene panels for colon cancer, prostate cancer, and several other cancer types are in development^20^.

Thus far, efforts to discover prognostic biomarkers have largely sought to identify gene expression changes associated with clinical outcome^21–24^. These studies have demonstrated that genes associated with cell cycle progression correlate with aggressive disease in multiple cancer types^21, 25–28^. Moreover, a series of recent reports have highlighted that copy number alterations (CNAs) also convey significant prognostic information, with increasing CNA burden generally associated with disease recurrence and metastatic dissemination^29–33^. However, existing studies suffer from several limitations: 1) Published analyses largely focus on single cancers and/or genomic data types (e.g., gene expression or DNA methylation or genetic mutations). A comprehensive comparison of prognostic biomarkers across both cancer types and genomic platforms has not been reported. 2) Biomarker research is potentially affected by the “file-drawer” problem, in which certain results (like the discovery of a new biomarker) are more likely to be published, while negative results may end up in a file drawer rather than in the academic literature^34–36^. Unbiased genome-wide biomarker studies can potentially counteract this publication bias and provide an accurate depiction of the prognostic landscape for a cancer of interest. 3) The Cancer Genome Atlas (TCGA), a project that collected genomic and clinical information from 33 cancer types^37^, has provided a rich resource for biomarker discovery. However, existing analyses^21, 38, 39^ - including our own prior work^29^ - have relied on preliminary survival data that was published on a cohort-by-cohort basis. A final set of updated and harmonized TCGA survival data, including the correction of more than 1,000 patients with annotation errors, has recently been published^40^, and these existing resources have not been revised.

Beyond the potential clinical relevance of prognostic biomarkers, molecular survival analysis has also become a staple of preclinical cancer research^41, 42^. With the increasing availability of genome-wide data from patient cohorts like TCGA^40^ and MSK-IMPACT^43^, it has become straightforward to assess whether a gene-of-interest is associated with patient outcome. These analyses typically seek to leverage survival data as clinical validation of the importance of a gene-of-interest in cancer biology. If the overexpression or mutation of some gene is associated with metastasis and patient death, then this is sometimes presented as evidence that that gene is a driver of cancer progression. Alternately, if the overexpression of a gene is associated with favorable prognosis, then it may be assumed that this gene has tumor-suppressive properties. Similar reasoning can underlie the prioritization of targets for therapeutic development: genes that are associated with poor prognosis are presumed to make the best drug targets due to their conjectured role as cancer drivers, while genes associated with favorable prognosis may be disregarded as non-essential for cancer progression^41^.

To our knowledge, the assumptions underlying these inferences have never been directly tested. While it seems intuitive that the presence of a genetic alteration that drives cancer progression would be associated with worse outcomes, it is not currently known whether real-world data supports this link. Moreover, the prognostic correlations of successful and unsuccessful cancer drug targets have not been investigated. In order to gain insight into the molecular differences between aggressive and indolent human cancers, and to critically evaluate the use of prognostic data in preclinical cancer research, we performed unbiased survival analysis from all cancer patients and all genomic data platforms included in TCGA.

## Results

### A comprehensive analysis of TCGA patient survival data

In order to identify the genomic features that correlate with cancer patient prognosis, we conducted a comprehensive analysis of outcome data for the 33 cancer types profiled by TCGA. Clinical endpoints were selected based on the updated data and recommendations provided in Liu et al.^40^: overall survival (OS) was used as an endpoint in 24 cancer types, while progression-free intervals (PFI) were used in 9 cancer types for which few deaths were observed during the study period (Table 1). For every patient cohort, we extracted information on six different features that were measured in individual tumors: point mutations, copy number alterations (CNAs), gene expression, microRNA expression, DNA methylation, and protein expression. For the mutational analysis, we considered only non-synonymous mutations, and in each patient cohort we excluded genes that were mutated in <2% of patients in that cohort (see the Materials and Methods). We then generated Cox proportional hazard models to assess the relationship between patient outcome and each individual gene for every tumor feature.

**Table 1.** Summary of the patient cohorts and data types included in this study.

| TCGA Cohort | Cancer type | Patient counts |  |  |  |  |  | Clinical endpoint |
| --- | --- | --- | --- | --- | --- | --- | --- | --- |
|  |  | Copy number alterations | DNA methylation | Gene expression | miRNA expression | Mutations | Protein expression |  |
| ACC | Adrenocortical carcinoma | 89 | 79 | 79 | 79 | 92 | 46 | Overall survival |
| BLCA | Bladder Urothelial Carcinoma | 404 | 411 | 404 | 405 | 406 | 340 | Overall survival |
| BRCA | Breast invasive carcinoma | 1063 | 1066 | 1089 | 1064 | 1015 | 882 | Progression-free interval |
| CESC | Cervical squamous cell carcinoma and endocervical adenocarcinoma | 294 | 306 | 304 | 306 | 289 | 173 | Overall survival |
| CHOL | Cholangiocarcinoma | 36 | 36 | 36 | 36 | 36 | 30 | Overall survival |
| COAD | Colon adenocarcinoma | 425 | 449 | 448 | 418 | 403 | 357 | Overall survival |
| DLBC | Lymphoid Neoplasm Diffuse Large B-cell Lymphoma | 48 | 48 | 48 | 47 | 37 | 33 | Progression-free interval |
| ESCA | Esophageal carcinoma | 182 | 183 | 184 | 182 | 184 | 126 | Overall survival |
| GBM | Glioblastoma multiforme | 571 | 428 | 155 | 0 | 390 | 232 | Overall survival |
| HNSC | Head and Neck squamous cell carcinoma | 516 | 522 | 519 | 518 | 507 | 212 | Overall survival |
| KICH | Kidney Chromophobe | 65 | 65 | 65 | 65 | 65 | 62 | Overall survival |
| KIRC | Kidney renal clear cell carcinoma | 506 | 521 | 533 | 499 | 369 | 478 | Overall survival |
| KIRP | Kidney renal papillary cell carcinoma | 282 | 285 | 289 | 285 | 280 | 214 | Overall survival |
| LAML | Acute Myeloid Leukemia | 179 | 178 | 161 | 175 | 130 | 0 | Overall survival |
| LGG | Brain Lower Grade Glioma | 509 | 512 | 514 | 508 | 509 | 427 | Progression-free interval |
| LIHC | Liver hepatocellular carcinoma | 365 | 372 | 369 | 367 | 361 | 183 | Overall survival |
| LUAD | Lung adenocarcinoma | 491 | 512 | 506 | 499 | 506 | 357 | Overall survival |
| LUSC | Lung squamous cell carcinoma | 481 | 484 | 496 | 461 | 479 | 323 | Overall survival |
| MESO | Mesothelioma | 86 | 86 | 86 | 86 | 81 | 62 | Overall survival |
| OV | Ovarian serous cystadenocarcinoma | 515 | 531 | 295 | 440 | 380 | 402 | Overall survival |
| PAAD | Pancreatic adenocarcinoma | 183 | 183 | 178 | 177 | 176 | 123 | Overall survival |
| PCPG | Pheochromocytoma and Paraganglioma | 161 | 178 | 179 | 178 | 178 | 79 | Progression-free interval |
| PRAD | Prostate adenocarcinoma | 489 | 495 | 497 | 491 | 494 | 352 | Progression-free interval |
| READ | Rectum adenocarcinoma | 154 | 155 | 159 | 152 | 149 | 130 | Progression-free interval |
| SARC | Sarcoma | 252 | 257 | 259 | 256 | 236 | 223 | Overall survival |
| SKCM | Skin Cutaneous Melanoma | 454 | 454 | 454 | 433 | 451 | 342 | Overall survival |
| STAD | Stomach adenocarcinoma | 433 | 435 | 409 | 428 | 432 | 351 | Overall survival |
| TGCT | Testicular Germ Cell Tumors | 133 | 133 | 134 | 133 | 129 | 104 | Progression-free interval |
| THCA | Thyroid carcinoma | 497 | 503 | 505 | 502 | 492 | 377 | Progression-free interval |
| THYM | Thymoma | 122 | 123 | 119 | 123 | 122 | 89 | Progression-free interval |
| UCEC | Uterine Corpus Endometrial Carcinoma | 518 | 534 | 531 | 522 | 529 | 438 | Overall survival |
| UCS | Uterine Carcinosarcoma | 56 | 57 | 57 | 56 | 57 | 48 | Overall survival |
| UVM | Uveal Melanoma | 80 | 80 | 80 | 80 | 80 | 12 | Overall survival |

To verify the overall fidelity of the clinical and genomic data, we conducted several control analyses. First, we examined survival curves for each of the 33 cancer types that comprise TCGA. Glioblastoma (GBM) exhibited the worst overall outcomes, with a median survival period of 428 days, while the pancreas adenocarcinoma (PAAD), acute myeloid leukemia (LAML), and mesothelioma (MESO) cohorts also exhibited very short survival times (Figure S1A). Consistent with previous reports, the 5-year survival frequencies for the TCGA cohorts were strongly correlated with national averages reported by NCI-SEER (Figure S1B; R = 0.83, P < .0001), suggesting that these patient cohorts are broadly representative of the general population^44^. Secondly, we confirmed that patient age, tumor stage, and tumor grade all exhibited a cancer type-independent association with shorter survival times, consistent with the well-established relationship between these clinical variables and poor outcomes (Figure S1C-E)^45, 46^. Third, we inferred chromosomal sex based on the expression of an X-chromosome marker (XIST) and a Y chromosome marker (RPS4Y1), and we found that the extrapolated values exhibited >99% concordance with the annotated gender of each patient (Figure S1F)^21^. Similarly, the methylation patterns of two X chromosome genes were also >99% concordant with gender (Figure S1G). Finally, we calculated the mutation frequencies of 106 verified oncogenes and tumor suppressors, and we found that these frequencies were highly similar to a previously reported pan-cancer analysis of TCGA data (Figure S1H; R = 0.99, P < .0001)^47^.

After establishing the validity of our analysis pipeline, we conducted two types of Cox analysis using the processed clinical and genomic data (Figure 1A). First, we generated univariate models, in which individual genomic features were directly associated with patient outcome (Table S1). Secondly, we generated multivariate (“fully-adjusted”) models, in which patient age, sex, tumor stage, and tumor grade were incorporated along with the genomic data (Table S2). For each Cox model, we report the Z score, which encodes both the directionality and significance of a survival relationship. Z scores across cancer types were combined using Stouffer’s method^48^. A Z score >1.96 indicates that the presence or upregulation of a tumor feature is associated with shorter survival times at a P < .05 threshold, while a Z score <-1.96 indicates that the presence or upregulation of a feature is associated with longer survival times at a P < .05 threshold. In general, the Z scores produced by the univariate and fully-adjusted models were highly concordant (R = 0.93-0.98), suggesting that few prognostic markers were affected by the inclusion of additional clinical variables (Figure S2). Because of this high degree of concordance, and in order to avoid the reported ambiguities in assessing clinical features like stage and grade^9–14^, we focus our analysis below on the genome-wide univariate models. Additionally, we created an online resource, available at http://survival.cshl.edu, to facilitate community access to this biomarker dataset.

**Figure 1.**
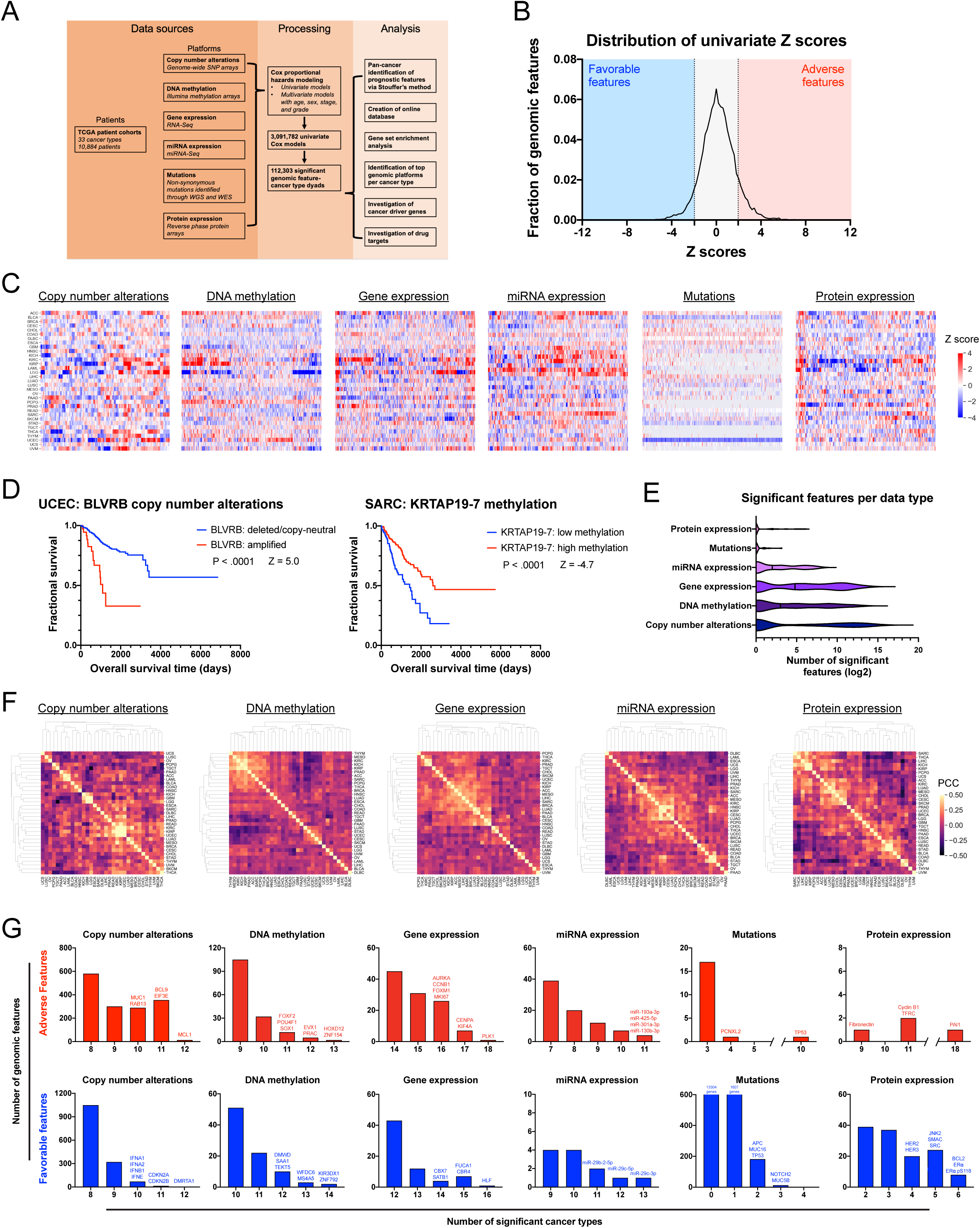
Pan-cancer, cross-platform identification of genomic features associated with patient outcome. (A) Schematic outline of the data processing and analysis performed for this work. (B) A density plot showing the distribution of genomic feature Z scores combined across all six platforms (CNAs, methylation, mutation, gene expression, miRNA expression, and protein expression). The dotted line at Z = −1.96 corresponds to P < .05 for a favorable feature, while the dotted line at Z = 1.96 corresponds to P < .05 for an adverse feature. (C) Heatmaps showing the distribution of Z scores within each of the six genomic platforms. Each row corresponds to a cancer type and each column corresponds to a gene or genomic feature. Note that, for Cox modeling of prognostic mutations, many genes were not mutated in ≥2% of patients in an individual cohort and so were not included in this analysis. The complete set of Z scores are included in Table S1. (D) Kaplan-Meier plots displaying two representative prognostic biomarkers identified from our genome-wide Cox modeling. Copy number amplification of BLVRB is associated with shorter survival times in UCEC (left). Methylation of KRTAP19-7 is associated with longer survival times in SARC (right). (E) Violin plots showing the distribution of significant prognostic features (|Z| > 1.96) per cancer type for each genomic platform. (F) Cluster plots of the correlation of Z scores for each genomic platform across cancer types. The scale indicates the strength of the Pearson correlation coefficient between Z score vectors. (G) Histograms displaying the number of shared prognostic biomarkers for each genomic platform across cancer types.

### Identification of genomic features that correlate with cancer patient prognosis

In total, we generated Z scores for 3,091,782 univariate Cox models (Figure 1A-C and Table S1). Across these models, we identified 112,303 genomic feature-cancer type dyads that were significantly associated with patient survival time at a Benjamini-Hochberg false-discovery rate of 1%. Two representative prognostic biomarkers discovered through this analysis are displayed in Figure 1D: CNAs affecting the gene encoding the heme metabolism enzyme BLVRB were identified as an adverse biomarker in endometrial carcinoma (UCEC), while methylation of the gene encoding the keratin-associated protein KRTAP19-7 was identified as a favorable biomarker in sarcoma (SARC). In general, gene expression, DNA methylation, and copy number alterations provided the most prognostic information, while mutational analysis provided the least (Figure 1E). Cancers that arose from related tissues of origin tended to display similar survival profiles (Figure 1F). For instance, Z scores derived from CNAs in low-grade glioma (LGG) were highly correlated with Z scores from GBM CNAs (R = 0.56, P < .0001), and Z scores produced from DNA methylation profiles were similar between renal clear cell carcinomas (KIRC) and renal papillary cell carcinomas (KIRP; R = 0.48, P < .0001).

By randomly permuting gene labels, we discovered that prognostic biomarkers were shared across multiple cancer types significantly more often than expected by chance (Figure S3). However, no single genomic feature was prognostic across all cancer types. The most broadly prognostic features were the expression of the RNA encoding the mitotic kinase PLK1 and the protein levels of the serine protease inhibitor PAI1, both of which were significantly associated with poor outcomes in 18 of 33 cancer types (Figure 1G). The most widely favorable biomarker was the expression of the transcription factor HLF, which was associated with better outcomes in 16 of 33 cancer types. Mutations in the tumor suppressor TP53 were an adverse event in 10 cancer types; no other mutation was associated with worse outcomes in more than four cancer types (Figure 1G and S4A). TP53 mutations were also found to correlate with longer survival times in GBM and lung squamous cell carcinomas (LUSC; Figure S4B). Mutations in TP53 have previously been recognized as a favorable prognostic biomarker in glioma while, to our knowledge, no such relationship has been observed in LUSC^49, 50^. Including TP53 mutations as a variable in multivariate Cox analysis did not significantly affect the identification of prognostic biomarkers (Figure S4C and Table S3).

Gene-level Z scores generated via RNA-Seq and protein-level Z scores generated via reverse-phase protein arrays (RPPA) were highly similar for features that were shared between these platforms (Figure S5). For instance, high expression of transferrin receptor (TFRC) mRNA and high expression of its protein product were both associated with poor outcomes in 11 different cancer types, including ACC (adrenocortical carcinoma) and LGG (Figure S5D). However, the same relationships were not apparent across all genomic platforms. Z scores from mutations, methylation profiling, and CNAs were only modestly correlated with Z scores at the transcript level (Figure S5A-C; R = −0.09 to R = 0.20). Methylation, mutations, and CNAs have complicated effects on downstream gene expression: for instance, inactivating mutations in TP53 can increase TP53 levels through a feed-forward mechanism^51, 52^, while certain chromosomal amplifications can fail to increase protein expression due to dosage compensation^53^. These results indicate that, while survival profiles generated at the level of transcription and protein expression are comparable, other classes of alterations can provide distinct prognostic information that is not captured solely by measuring the expression of the affected gene(s).

### Identification of gene sets and pathways broadly associated with cancer patient outcome

We next sought to understand the biological pathways that were differentially-regulated in deadly cancers. We performed gene ontology (GO) enrichment analysis on the genes that were identified by RNA-Seq as cross-cancer prognostic biomarkers at a Benjamini-Hochberg false-discovery rate of 1%. Consistent with previous results^21, 28, 54^, we observed that transcripts over-expressed in aggressive tumors were highly-enriched for genes associated with chromosome segregation, DNA replication, and the mitotic cell cycle (Figure 2A-C, Table S1C and S4A). This gene set included several known proliferation markers (MCM2, MKI67, PCNA)^55^, and promoter analysis revealed that many adverse genes were controlled by the cell cycle-regulated E2F family of transcription factors (Figure 2B and Figure S6A-B)^56^. Within the protein expression dataset, cell cycle-associated proteins including Cyclin B1 and MSH6 were also among the top-scoring adverse features (Figure S6C and Table S1F). Finally, we observed that the expression of the adverse transcripts was tightly correlated with both cancer cell line doubling times measured *in vitro* and a direct analysis of mitotic activity in tumor specimens (Figure 2D and Figure S6D>)^57^. In total, these results suggest that the expression of these adverse biomarkers reflect a tumor’s proliferative rate. In contrast, favorable expression markers included ACAD11, INPP5K, and CHP, and were enriched for genes whose products localize to mitochondria and are involved in catabolic processes (Table S1C and S4B).

**Figure 2.**
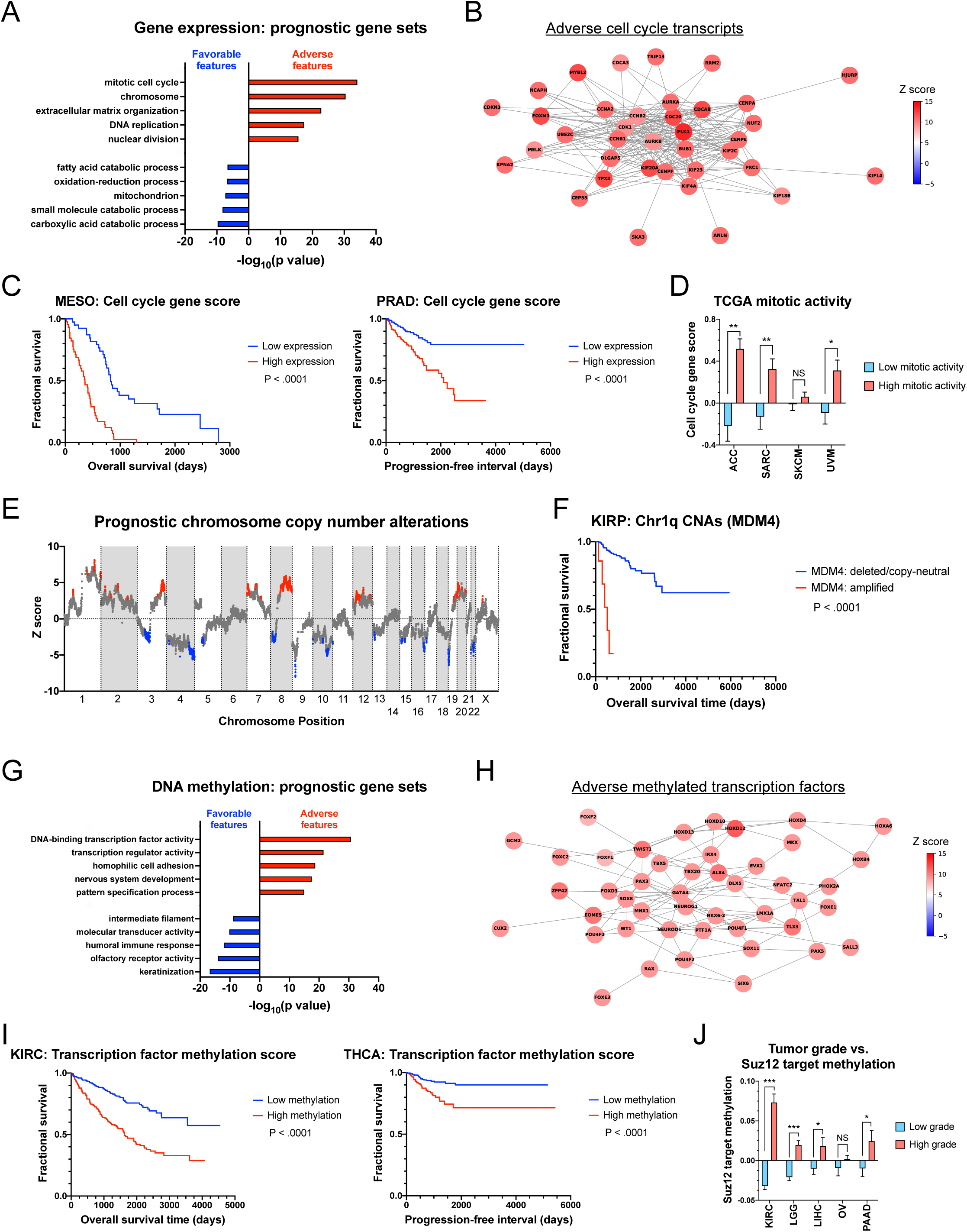
Identification of prognostic gene sets across genomic platforms. (A) GO terms enriched among adverse and favorable gene expression biomarkers. The complete set of GO terms are included in Table S4A-B. (B) Network of interactions among cell cycle genes, colored according to Stouffer’s Z from the combined gene expression Cox models. (C) Kaplan-Meier plots displaying survival in MESO (left) and PRAD (right) based on the mean expression of a set of transcripts associated with the gene ontology term “mitotic cell cycle”. (D) Bar graph showing cell cycle scores based on pathologically-observed mitotic activity in different TCGA cohorts. (E) A plot displaying Stouffer’s Z by chromosomal coordinate. Red dots indicate loci where genomic amplifications are associated with worse outcomes and blue dots indicate loci where genomic deletions are associated with worse outcomes. The complete list of genes found within these regions is included in Table S5. (F) Kaplan-Meier plot displaying survival in KIRP split based on the copy number status of the Chr1q gene MDM4. (G) GO terms enriched among adverse and favorable methylation biomarkers. The complete set of GO terms are included in Table S4E-S4F. (H) Network of interactions among developmental transcription factors colored according to Stouffer’s Z from the combined methylation Cox models. (I) Kaplan-Meier plots displaying survival in KIRC (left) and THCA (right) based on the methylation of a collection of developmental transcription factors. (J) Bar graph showing the average methylation of Suz12 targets based on tumor grade in different TCGA cohorts.

To find genomic loci where CNAs were associated with patient outcome, we generated a profile of Z scores by chromosomal coordinates (Figure 2E). We applied a peak-finding algorithm to identify genomic “peaks”, which correspond to loci where copy number gains are associated with poor outcomes, and “valleys”, which correspond to loci where deletions are associated with poor outcomes. GO term enrichment analysis revealed that very few gene sets were significantly enriched in either peaks or valleys (Table S4C-D). This indicates that CNAs affecting many different cellular pathways are enriched in deadly tumors. However, manual inspection revealed that a number of known oncogenes and tumor suppressors are encoded in these regions. For instance, the highest peak is found on the q arm of chromosome 1 and encompasses the known cancer driver gene MDM4 (Figure 2F). The deepest valley is found on Chr9p and is centered on the cell cycle inhibitor CDKN2A, deletion of which has previously been associated with deadly cancers^29, 58^. Finally, we determined prognostic peaks and valleys from CNA data for each of the 33 individual cancer types (Table S5). We speculate that some of these regions may harbor genes with uncharacterized roles in cancer biology.

Gene ontology analysis of methylation events in tumors with grim prognosis revealed a striking enrichment of transcription factors and genes involved in embryonic development, including *NKX6-1*, *HOXD12*, and *FOXE1* (Figure 2G-I, Table S1B and S4E). Favorable methylation events were more diverse and included genes encoding intermediate filaments, olfactory receptors, and keratin-associated proteins (Table S4F). Certain cancers exhibit *de novo* methylation of developmental genes that are silenced by the chromatin-modifying Polycomb complex during embryogenesis^59, 60^. These Polycomb targets include lineage-defining transcription factors that are activated or repressed to specify tissue identity. Our finding that developmental transcription factors were methylated in aggressive tumors led us to investigate whether these adverse features were also linked with Polycomb activity. Indeed, we observed a highly-significant enrichment of Polycomb component Suz12 binding sites among the genes where high levels of methylation were associated with shorter patient survival (Figure S8A)^61^. In contrast, Suz12 sites were under-represented among genes where high levels of methylation were favorable features. Expression of EZH2, which encodes the catalytic subunit of the Polycomb complex, and the DNA methyltransferases DNMT1, DNMT3A, and DNMT3B, which cooperate with Polycomb to silence target loci^62^, were all identified as significant pan-cancer adverse features (Figure S8B-C and Table S1A). While the genome-wide correlation between methylation and transcription-associated survival profiles was minimal (Figure S5A), we found that methylation of these Polycomb-associated loci in particular was associated with decreased gene expression (Figure S8D). Finally, we observed that methylation of these adverse biomarkers was frequently observed in high-grade (dedifferentiated) malignancies (Figure 2J). This data suggests that cancers methylate and silence lineage-defining transcription factors, which promotes the loss of cell identity and is associated with aggressive disease.

### Cross-platform identification of the most informative prognostic biomarkers per cancer type

Given the ability to interrogate any gene on any genomic platform in a primary tumor, what measurements confer the most prognostic information? To address this question, we identified the 100 features in each cancer type that exhibit the strongest overall correlations with patient outcome in both univariate and fully-adjusted models (Figure 3A). We found that, for the average cancer type, 46% of top-scoring features were gene expression measurements, 22% were methylation events, 30% were copy number alterations, and only 1% were mutations. Some cancers diverged from this overall trend: in GBM, 95 of the top 100 univariate biomarkers were favorable methylation events, which likely reflects the CpG island methylator phenotype that has been linked with long-term survival in this cancer type^63^. Interestingly, several top prognostic features within individual cancer types are poorly-characterized. For instance, in stomach adenocarcinoma (STAD), across 96,730 Cox models that we generated, the genomic feature that exhibited the strongest association with patient outcome was expression of the uncharacterized lncRNA FLJ16779/LOC100192386. Finally, we found that classifying patients based on single features identified through this analysis was sufficient to distinguish outcomes in ambiguous clinical situations where patients are at risk of undertreatment or overtreatment. This includes stage 1A breast cancer^64^, stage 2 colon cancer^65, 66^, and Gleason 7 prostate cancer^67, 68^ (Figure 3B).

**Figure 3.**
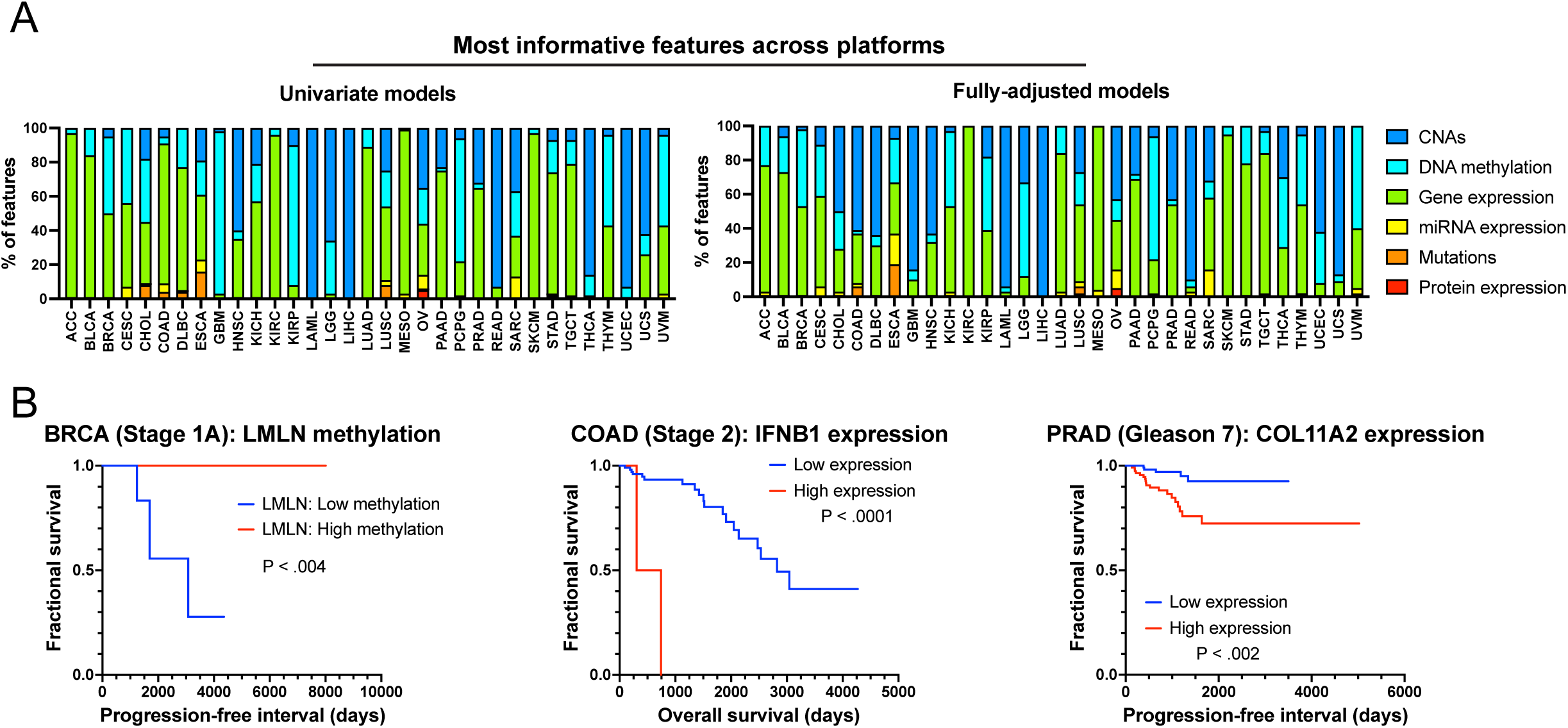
Cross-platform analysis reveals that cancer gene expression measurements harbor the most prognostic information. (A) The 100 genomic features that exhibit the strongest associations with patient outcomes in univariate or fully-adjusted Cox models are displayed. (B) Kaplan-Meier plots displaying patient survival in stage 1A breast cancer, stage 2 colon cancer, and Gleason 7 prostate cancer, split based on the indicated biomarkers.

### Over-expression or mutation of verified cancer driver genes is not widely associated with poor prognosis

When the expression or mutation of a gene is found to be associated with poor patient prognosis, this is typically presented as evidence that that gene is an important driver of disease progression^41^. However, we were surprised to find that very few established oncogenes or tumor suppressors were identified as significant adverse features in our genome-wide analyses described above. Commonly-mutated cancer driver genes including KRAS, HRAS, PIK3CA, CTNNB1, RB1, and APC were not recovered as prominent biomarkers. Instead, the strongest biomarkers in our expression analysis tended to be housekeeping genes with roles in the cell cycle. In our sequencing analysis, TP53 mutations were identified as an adverse feature in 10 of 33 cancer types, but no other gene was significant in more than four cancer types. These findings challenged the notion that we should expect important cancer driver genes to be associated with patient outcomes. We therefore decided to study these unexpected results more closely.

To systematically examine the prognostic significance of mutations in cancer driver genes, we assessed two collections of verified oncogenes. First, we considered a set of 31 genes that exhibit pan-cancer oncogenic activity (BRAF, EGFR, KRAS, etc.)^47^, and we calculated Z scores for these genes in each of the 33 TCGA cancer types. Secondly, we considered an expanded set of 81 oncogenes, but we only calculated Z scores for these genes in cancer types in which that gene is recurrently activated (e.g., FLT3 in LAML, EGFR in LUAD, ERBB2 in BRCA, etc.)^47^. We found that, considered as a group, the Z scores for these oncogenes were not enriched for prognostic features relative to randomly-permuted gene sets of the same size (Figure 4A-B). The mean Z score for the pan-cancer oncogene set was −0.06, and the mean Z score for the tissue-limited oncogene set was −0.08. Out of 1023 possible cancer type-oncogene pairs, we found that mutations in a pan-cancer oncogene were associated with worse prognosis in <1% of instances, which was not significantly different from the background rate of prognostic mutations across all genes (Figure 4C). Analyzing individual Kaplan-Meier curves supported these findings. For example, EGFR is a known driver of glioblastoma^69^, but EGFR mutations were not associated with poor prognosis in the TCGA GBM cohort (Figure 4D). Indeed, in several cases, we observed that mutations in established driver oncogenes like CTNNB1 and ERBB4 were associated with favorable outcomes relative to cancers that lacked mutations in these genes (Figure 4E).

**Figure 4.**
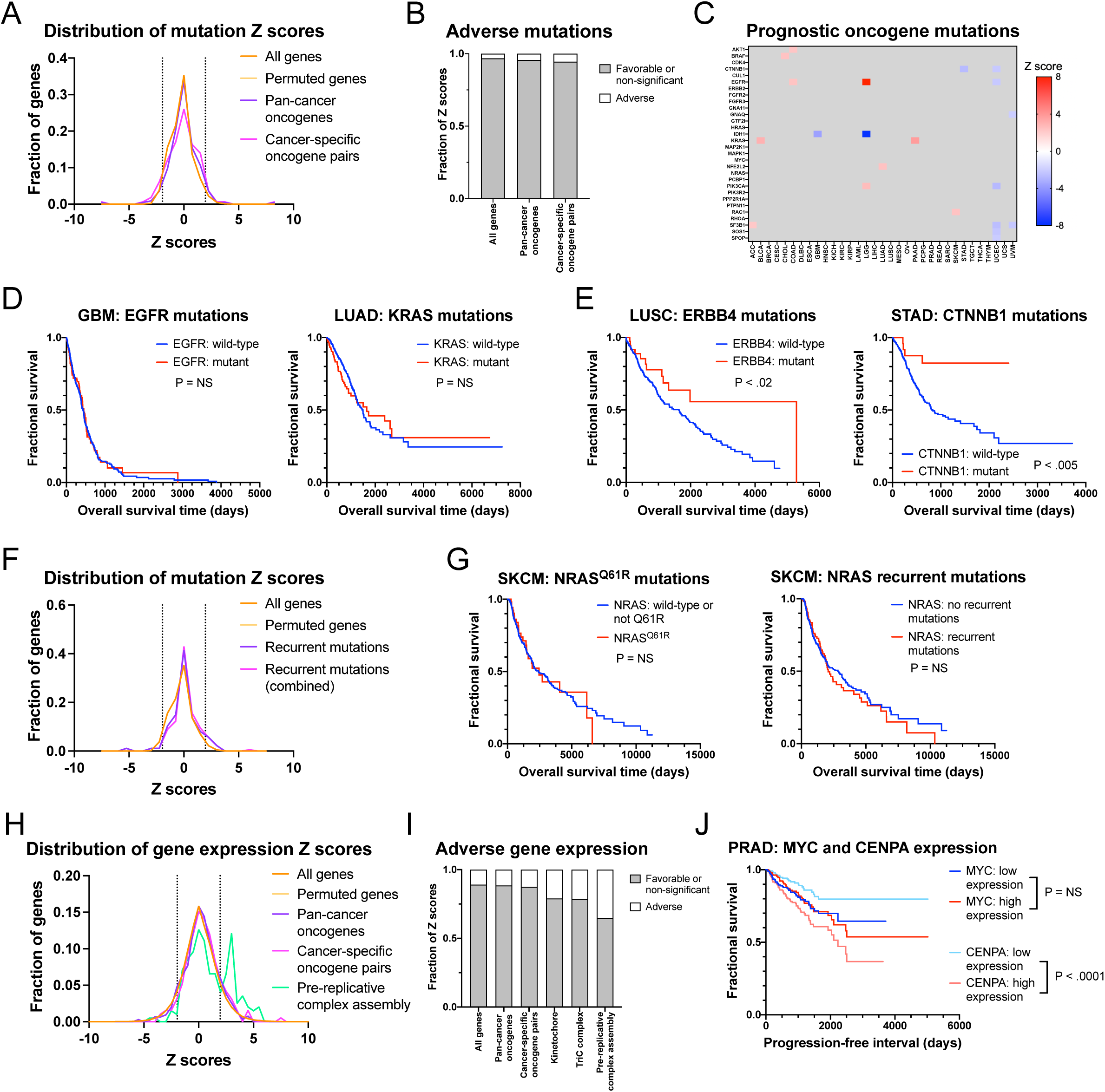
Oncogene mutation or overexpression is not widely associated with patient outcome. (A) A density plot showing the distribution of mutation Z scores for the indicated gene sets. The dotted line at Z = −1.96 corresponds to P < .05 for a favorable mutation, while the dotted line at Z = 1.96 corresponds to P < .05 for an adverse mutation. (B) Stacked bar graph showing the fraction of mutations that are associated with adverse outcomes for the indicated gene sets (C) A heatmap showing significant (|Z| > 1.96) survival associations for oncogene mutations. Each row represents a pan-cancer oncogene (identified in Bailey et al.^47^) and each column represents a patient cohort from TCGA. (D) Kaplan-Meier plots showing that mutations in the established GBM oncogene EGFR (left) and the established LUAD oncogene KRAS (right) are not associated with shorter survival times. (E) Kaplan-Meier plots showing that mutations in the LUSC oncogene ERBB4 (left) and the STAD oncogene CTNNB1 (right) are associated with longer survival times. (F) A density plot showing the distribution of mutation Z scores for the indicated gene sets, including “hotspot” mutations that affect specific recurrently-mutated codons. The dotted line at Z = −1.96 corresponds to P < .05 for a favorable mutation, while the dotted line at Z = 1.96 corresponds to P < .05 for an adverse mutation. The complete list of mutation Z scores are included in Table S6. (G) Kaplan-Meier plots demonstrating that the most common NRAS driver mutation - Q61R - or a combination of all common NRAS driver mutations - G12D, G12S, G13D, G13R, K16N, Q61H, Q61K, Q61R, and E62K - are not associated with shorter survival times in SKCM. (H) A density plot showing the distribution of gene expression Z scores for the indicated gene sets. Note that while the oncogene sets largely overlap with the control gene sets, the pre-replicative complex assembly gene set displays a significant peak to the right of the control gene sets. The dotted line at Z = −1.96 corresponds to P < .05 for favorable overexpression, while the dotted line at Z = 1.96 corresponds to P < .05 for adverse overexpression. (I) Stacked bar graph showing the fraction of gene expression biomarkers that are associated with adverse outcomes for the indicated gene sets. (J) Kaplan-Meier plot showing survival in PRAD split based on the expression of a key driver oncogene (MYC) or split based on the expression of a non-oncogenic cell cycle gene (CENPA).

In the above analysis, all cancers that harbored a non-synonymous mutation in a gene of interest were classified as “mutant” for that gene. While we recognize that different mutations may have different functional effects, manual inspection of the data revealed that most of these mutations were likely to exhibit oncogenic activity. For instance, >95% of non-synonymous mutations in KRAS were found in codons 12, 13, or 61, which have all been identified as sites of driver mutations^70^. However, we considered the possibility that our above analysis could be obscuring certain specific mutations with prognostic significance. Accordingly, we conducted two further analyses: 1) we analyzed individual recurrent mutations separately (e.g., we calculated Z scores separately for tumors with KRAS^G12V^ mutations, KRAS^G12R^ mutations, KRAS^G13D^ mutations, etc.) and 2) we combined patients with any recurrently-observed mutation within a gene but excluded patients with non-synonymous mutations that were not found in a commonly-altered codon. However, these analyses were largely consistent with our whole-gene analysis and revealed few prognostic mutations (Figure 4F, S8, and Table S6). Many of the most common cancer driver mutations, including PIK3CA^E545K^, KRAS^G13D^, IDH1^R132H^, and FBXW7^R465H^, were not associated with worse outcomes in any of the 33 TCGA cohorts (Figure S8). For instance, while NRAS mutations are an established driver of melanoma, neither the most common NRAS alteration (Q61R) nor a combination of all common NRAS alterations predicted poor survival (Figure 4G)^71^. In total, these results indicate that mutations in verified cancer driver genes like KRAS, PIK3CA, and IDH1 are not widely associated with adverse outcomes.

Next, we investigated whether the over-expression of verified oncogenes was associated with shorter survival times. We calculated Z scores for the two collections of driver oncogenes described above and compared them to randomly-permuted gene sets of the same size. Consistent with our mutational analysis, we found that the expression of verified oncogenes was no more likely to be an adverse prognostic feature than the expression of a randomly-selected gene (Figure 4H). Indeed, we found that oncogenes harbored less prognostic power than sets of genes encoding the kinetochore, the TriC complex, or the pre-replication complex, which are not known to harbor oncogenic activity but are associated with cell cycle progression (Figure 4I). For instance, in prostate cancer (PRAD), we observed that high expression of the driver oncogene MYC was not associated with adverse outcomes, but high expression of the kinetochore component CENPA was strongly associated with disease progression (Figure 4J). Taken together, these results demonstrate that the expression and/or mutation of key cancer driver genes is not a robust predictor of poor patient outcomes.

### Successful cancer drugs generally do not target adverse prognostic genes

Many papers characterizing a novel drug or drug target in cancer biology present evidence that the overexpression or mutation of that drug target is associated with aggressive disease^41^. The assumption underlying this line of evidence is that genes that are associated with shorter survival times make the best targets for therapeutic development. However, to our knowledge, this assumption has never been directly tested. We therefore set out to explore whether successful cancer drugs that currently exist are likely to target genes that are associated with poor prognosis.

We generated a list of FDA-approved cancer drugs matched with each drug’s reported target(s) (Table S7A)^72^. We then calculated Z scores for each drug target in the cancer type(s) for which that drug has received FDA approval. Surprisingly, we found that successful drug targets were not generally enriched for adverse prognostic factors (Figure 5A-D and Table S7B-C). Out of 212 target-cancer type pairs, mutation of <2% of targets was associated with worse outcomes. Among gene expression biomarkers, drug target Z scores were not significantly greater than the Z scores of randomly-permuted gene sets. In fact, we observed that FDA-approved drugs were as likely to target a gene whose expression correlated with favorable prognosis as they were to target a gene whose expression correlated with poor prognosis (12% vs. 17%, P = NS; Figure 5D).

**Figure 5.**
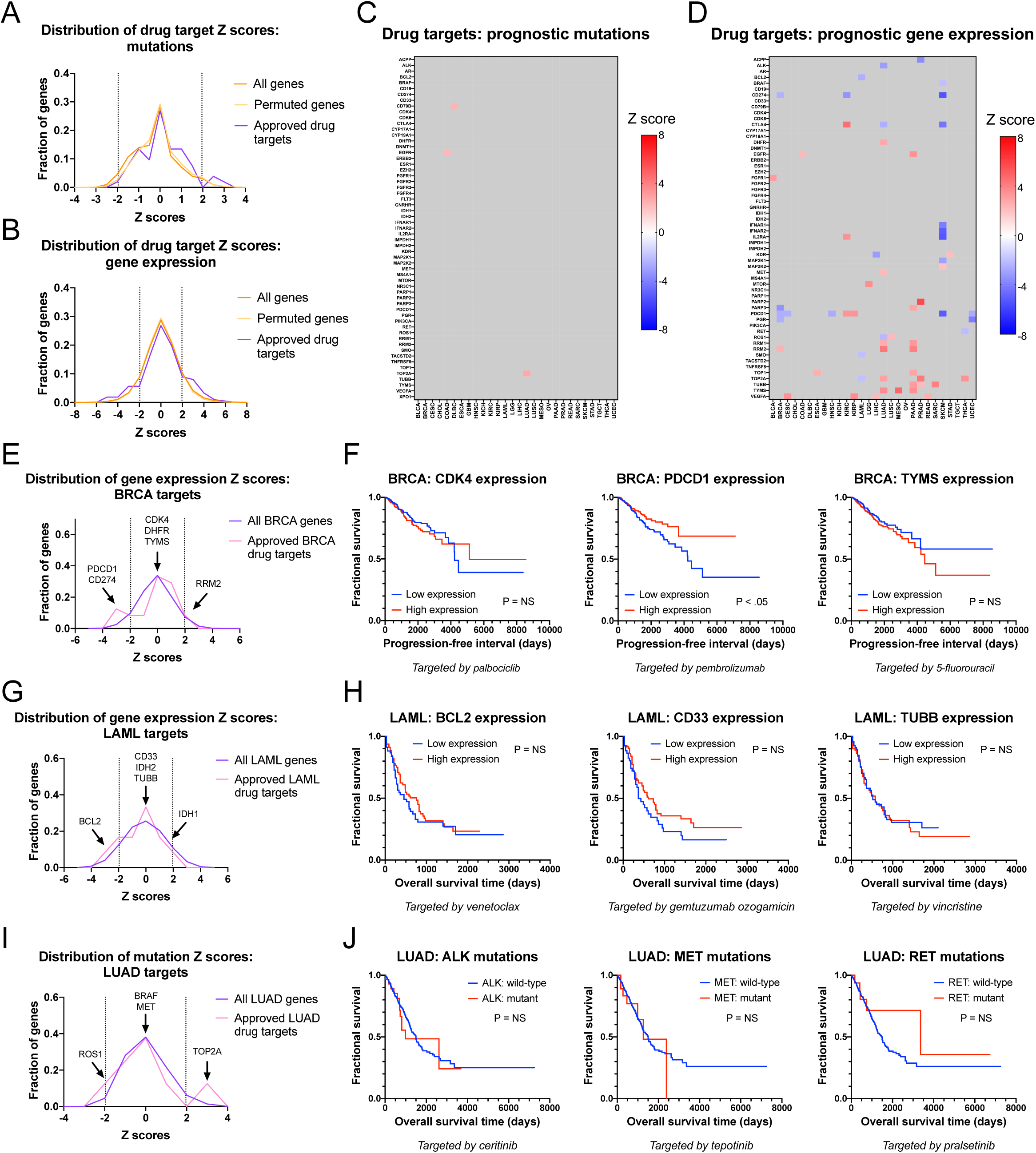
The genomic targets of FDA-approved cancer drugs are not strong prognostic biomarkers. (A) A density plot showing the distribution of mutation Z scores for the indicated gene sets. The dotted line at Z = −1.96 corresponds to P < .05 for a favorable mutation, while the dotted line at Z = 1.96 corresponds to P < .05 for an adverse mutation. (B) A density plot showing the distribution of gene expression Z scores for the indicated gene sets. The dotted line at Z = −1.96 corresponds to P < .05 for favorable overexpression, while the dotted line at Z = 1.96 corresponds to P < .05 for adverse overexpression (C) A heatmap showing significant (|Z| > 1.96) survival associations for mutations in the targets of FDA-approved drugs. Each row represents a drug target and each column represents a cancer patient cohort from TCGA. (D) A heatmap showing significant (|Z| > 1.96) survival associations for expression changes in the targets of FDA-approved drugs. Each row represents a drug target and each column represents a cancer patient cohort from TCGA (E) A density plot showing the distribution of gene expression Z scores for the indicated gene sets in BRCA. The dotted line at Z = −1.96 corresponds to P < .05 for favorable overexpression, while the dotted line at Z = 1.96 corresponds to P < .05 for adverse overexpression. (F) Kaplan-Meier plots displaying survival times in BRCA based on the expression levels of the BRCA drug targets CDK4, PCDC1, and TYMS. (G) A density plot showing the distribution of gene expression Z scores for the indicated gene sets in LAML. The dotted line at Z = −1.96 corresponds to P < .05 for favorable overexpression, while the dotted line at Z = 1.96 corresponds to P < .05 for adverse overexpression. (H) Kaplan-Meier plots displaying survival times in LAML based on the expression levels of the LAML drug targets BCL2, CD33, and TUBB. (I) A density plot showing the distribution of mutation Z scores for the indicated gene sets in LUAD. The dotted line at Z = −1.96 corresponds to P < .05 for favorable mutations, while the dotted line at Z = 1.96 corresponds to P < .05 for adverse mutations. (J) Kaplan-Meier plots displaying survival times in LUAD based on the mutations in the indicated LUAD drug targets ALK, MET, and RET.

Closer inspections of individual genes and patient cohorts revealed multiple factors that contribute to the minimal overlap between adverse biomarkers and successful drug targets (Figure 5E-J). First, we observed that some genes were strongly upregulated in certain cancer types, but within those cancer types, expression or mutation of that gene was non-prognostic. For instance, the FDA-approved LAML therapy gemtuzumab ozogamicin consists of a DNA damaging agent conjugated to an antibody targeting the CD33 antigen. CD33 expression is strongly upregulated in myeloid cells^73^, which confers specificity to this agent, but within LAML, variation in CD33 levels do not correlate with aggressive disease (Figure 5H). Secondly, many targetable driver mutations are mutually-exclusive and serve to activate the same signaling pathway^74–76^. For instance, lung cancers can harbor targetable mutations in ALK, BRAF, EGFR, MEK1, MET, RET, or ROS1, each of which activates the MAPK signaling pathway. There is no *prima facie* reason to believe that any one of these genes would be associated with worse prognosis than all others, and in the LUAD dataset, none of these mutations were correlated with outcome (Figure 5J and Table S1E).

Third, many cancer therapies exhibit significant cancer cell non-autonomous effects. For instance, breast tumors with high expression levels of PDCD1 (PD1) have superior outcomes relative to breast tumors with low PDCD1 expression (Figure 5F). Based on this survival correlation, one might assume that inhibiting PDCD1 would *decrease* patient survival. However, antibodies like pembrolizumab that inhibit PD1 have a pronounced benefit in multiple cancer types, including breast cancer^77–79^. In this case, PD1 is expressed by infiltrating immune cells^80, 81^, and strong immune infiltration enhances tumor control, even in the absence of immune checkpoint inhibitor treatment^82^. Finally, some targetable genes play key roles in cancer biology, even though their expression levels are uncorrelated with disease severity. For instance, thymidylate synthetase (TYMS) is required for DNA replication, and TYMS inhibitors like 5-fluorouracil are effective at blocking DNA replication in several cancer types^83, 84^. TYMS inhibitors can thereby prolong patient survival, even though TYMS is not known to be an oncogene and TYMS upregulation does not drive disease progression (Figure 5F). In total, these and other factors may contribute to our observation that a large majority of successful cancer drugs do not target genes that are associated with poor patient outcomes.

We also considered an alternate explanation for these results: it could be the case that the mutation or overexpression of a drug target renders tumors sensitive to that therapy, and so cancers expressing a druggable marker may have a favorable prognosis compared to cancers with an unknown or undruggable marker. If this were the case, then our discovery that few successful cancer drugs target prognostic factors could be a reflection of treatment, rather than the underlying cancer biology. To investigate this possibility, we conducted an additional analysis using only drug/indication pairs that received FDA approval in 2018 or later. As TCGA patient follow-up stopped in 2015/2016, we expect that extremely few patients in these cohorts would have received these therapies. However, our findings with this subset of drugs were consistent with our analysis of the complete dataset and revealed very few prognostic correlations among drug targets (Figure S9A-B). For instance, the FGFR inhibitor erdafitinib received FDA approval for use in FGFR3-mutant bladder cancer in 2019^85^, but neither FGFR3 mutations nor FGFR3 overexpression were prognostic in the TCGA BLCA cohort collected prior to this time (Figure S9C-D). In total, these results demonstrate that successful cancer drugs generally do not target biomarkers associated with aggressive disease.

### Many therapies targeting adverse prognostic factors have failed in clinical trials

To further evaluate the relative importance of targeting genetic features that are associated with aggressive tumors, we focused on the 50 genes whose expression exhibits the strongest correlation with adverse outcomes across cancer types (Table S1C). Using www.clinicaltrials.gov and other related resources, we identified therapeutic agents designed to target these top-scoring genes that have been tested in patients (Figure 6A-B). We found that 16 of the top 50 genes have been targeted in clinical trials, but therapies against 15 of these genes have failed to receive FDA approval. For instance, the top-scoring prognostic factor in our analysis is the mitotic kinase PLK1, and small-molecule compounds designed to block PLK1 activity have been tested in patients with multiple cancer types^86^. However, PLK1 inhibitors were found to cause severe and sometimes fatal side effects, thwarting their clinical utility^87^. Other inhibitors against top-scoring genes, including CDK1, AURKA, AURKB, and CENPE, were similarly found to exhibit unacceptable toxicities or insufficient activity (Figure 6A). Peptides derived from several top-scoring genes have also been used in immunotherapy vaccines, though their therapeutic efficacy has not been demonstrated in randomized trials. Of the 50 genes that exhibit the strongest correlations with aggressive disease, only a single gene, RRM2, is targeted by an FDA-approved compound. RRM2 encodes a subunit of ribonucleotide reductase, and ribonucleotide reductase inhibitors including hydroxyurea and gemcitabine have received FDA approval for use in several cancer types^88^.

**Figure 6.**
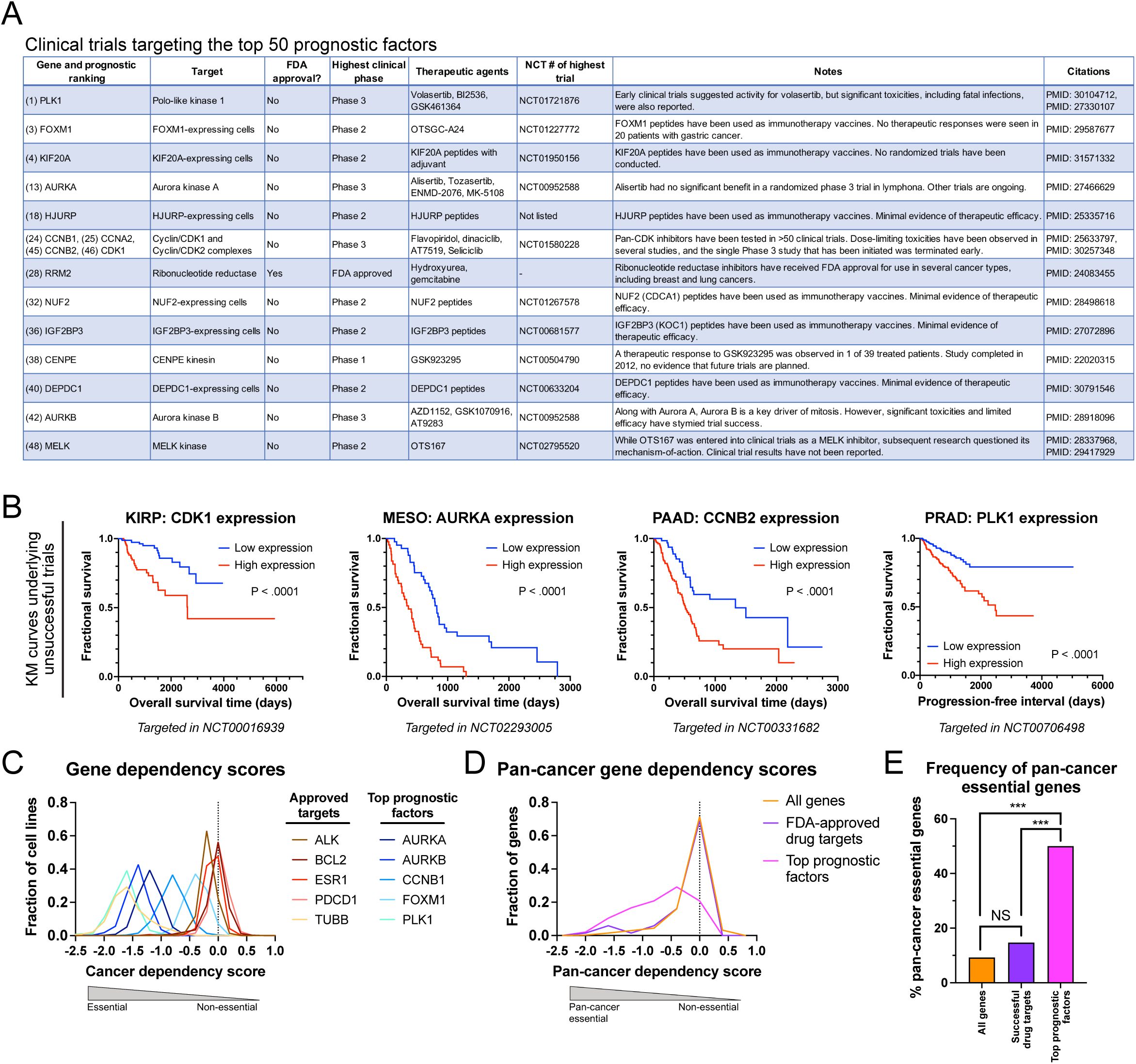
Therapies targeting top prognostic genes have failed in clinical trials. (A) A table displaying the genes among the 50 prognostic factors that exhibit the strongest correlations with cancer patient outcomes that have been targeted in cancer clinical trials. (B) Kaplan-Meier plots showing patient survival in the indicated cancer cohorts. Each graph displays a gene that has been targeted in clinical trials in that cancer type. (C) A density plot showing the distribution of cancer dependency scores for the indicated genes, split according to whether the gene is the target of an FDA-approved cancer therapy or whether the gene is a top-scoring prognostic factor. (D) A density plot showing the distribution of pan-cancer cancer dependency scores for the indicated gene sets. (E) A bar graph showing the percent of genes that are essential across cancer types in the indicated gene sets.

In our gene ontology analysis, we noted that many top-scoring prognostic factors were widely-expressed housekeeping genes with crucial roles in cell cycle progression. We hypothesized that the frequent failure of these top-scoring genes as cancer drug targets could result from the fact that many of them represent broadly essential genes, potentially leading to significant side effects when their proteins are inhibited. To investigate this hypothesis, we analyzed cancer dependency scores from whole-genome CRISPR screening data across several hundred cancer cell lines^89, 90^. Each score measures the fitness effects of ablating the gene in question, with larger negative scores indicating more significant fitness defects upon gene loss. We observed that the dependency score distribution for FDA-approved cancer targets was very similar to the distribution of scores across all genes (Figure 6C-D). While some approved drugs inhibit pan-cancer dependencies (e.g., TUBB, TOP1, TOP2), a majority of targeted genes exhibit more selective effects (e.g., ESR1, PARP1, ALK). In contrast, the top-scoring prognostic factors exhibit essentiality patterns that are significantly different from the essentiality patterns of approved drugs. We observed that 50% of top-scoring prognostic factors are essential across all cell types, compared to only 15% of genes targeted by approved drugs (Figure 6E). This limited cell type-selectivity could contribute to the toxicity and high failure rate of drugs designed to target these top-scoring genes. In total, these analyses suggest that prioritizing targets for therapeutic development based on prognostic correlations may be counterproductive, as a large fraction of these correlated factors represent ubiquitously-expressed housekeeping genes rather than cancer-specific dependencies.

## Discussion

Genomic analysis has the potential to shed unprecedented insight into the molecular architecture of human cancers. In light of studies demonstrating both the pervasive over-treatment and under-treatment of cancer patients, the use of genomic technologies to discover and validate prognostic biomarkers could greatly enhance risk prediction and clinical treatment decisions. In this work, we generated a rich dataset of more than 3,000,000 individual Cox proportional hazards models and identified more than 100,000 significant prognostic biomarkers across 33 cancer types. These data have also been shared via a web portal at http://survival.cshl.edu to facilitate further analysis with this resource.

Our study illustrates the unexpected prognostic potential of different classes of genomic data. For instance, while there has been substantial attention devoted toward developing routine whole-exome and targeted sequencing panels for clinical use^18, 91^, our findings demonstrate that relatively few point mutations are significantly associated with cancer patient outcome. Aside from mutations in TP53, which were prognostic in 12 of 33 patient cohorts, mutations in established cancer genes like CDKN2A, EGFR, KRAS, PIK3CA, PTEN, RB1, and many others had extremely limited prognostic power. Sequencing oncogenes and tumor suppressors may be useful in order to assign patients to specific targeted therapy regimens, and larger patient cohorts sequenced at greater depths may identify prognostic relationships not found in this study. Nonetheless, considered as a whole, this work suggests that routine sequencing of patient tumors will not yield significant improvements in risk prediction relative to other potential genomic platforms.

In contrast to the paucity of prognostic mutations uncovered through this work, we identified several hundred genes whose methylation was associated with outcome across cancer types. Gene ontology analysis demonstrated that these genes were enriched for developmental transcription factors, and these genes were typically downregulated in high-grade tumors. Polycomb activity may thereby facilitate cancer cell reprogramming and a loss of cellular identity, returning cancers to a stem cell-like state that can rapidly progress^60^. Similarly, the most penetrant gene expression biomarkers were cell cycle-associated transcripts that were upregulated in cancers with high mitotic activity. Importantly, cellular proliferation has a profound influence on gene expression genome-wide, and so many genes with diverse functions may still be prognostic by indirectly capturing cell cycle activity^28^. In future work, the construction of multivariate Cox models incorporating proliferation markers like MKI67 and PCNA may help differentiate between cell cycle-dependent and cell cycle-independent prognostic features.

Our findings also have significant implications for the analysis of cancer survival data in a preclinical or therapeutic-discovery setting. Using multiple datasets of verified oncogenes, we unambiguously demonstrate that genes that drive tumorigenesis are not significantly enriched among adverse biomarkers within cohorts of cancer patients. Similarly, while genes associated with metastasis and patient death are sometimes presumed to encode the most promising targets for therapeutic development, we show that successful cancer drugs generally do not target adverse biomarkers. Correspondingly, a large majority of experimental drugs that do target adverse biomarkers have failed in clinical trials. We believe that these results underscore a crucial distinction between causation and correlation in clinical observations. To illustrate, among a random group of adults, individuals receiving kidney dialysis are more likely to die than individuals who are not receiving dialysis. Based strictly on this correlative observation, one could assume that kidney dialysis kills people. Yet, we know that people receiving dialysis are likely to be older and have several medical comorbidities, and dialysis saves their lives^41, 92^.

In general, we caution that deducing functional relationships and prioritizing drug targets based on cancer survival data may be inappropriate, and that such relationships may be fraught with confounding variables and spurious correlations. From our analysis, one could incorrectly infer that KRAS mutations are not important in lung cancer (Figure 4D), or that kinetochore gene expression is a more important driver of prostate cancer than MYC expression (Figure 4J). Moreover, we observed that strongly-prognostic genes tend to be widely essential across cell types, which could explain why so many therapies designed against these genes have exhibited dangerous side effects in human patients. Consequently, leveraging survival analysis to select targets for therapeutic development could inadvertently prioritize targets that are unlikely to succeed in clinical testing. Functional studies in which causative relationships can be interrogated are necessary to rigorously identify potential drug targets and genes that drive cancer progression. We suggest that, in general, the use of survival data to identify prognostic biomarkers should be decoupled from the use of survival data to infer gene function in cancer biology.

### Materials and Methods Data sources

TCGA data was acquired from the TCGA PanCanAtlas^93^. Final datasets used for this analysis include:

DNA copy number: *broad.mit.edu_PANCAN_Genome_Wide_SNP_6_whitelisted.seg*

DNA methylation: *usc.edu_PANCAN_merged_HumanMethylation27_HumanMethylation450. betaValue_whitelisted.tsv*

Gene expression: *EBPlusPlusAdjustPANCAN_IlluminaHiSeq_RNASeqV2.geneExp.tsv*

miRNA expression: *pancanMiRs_EBadjOnProtocolPlatformWithoutRepsWithUnCorrectMiRs*_08_04_16.csv

Mutations: *mc3.v0.2.8.PUBLIC.maf.gz*

Protein expression: *TCGA-RPPA-pancan-clean.txt*

For all cancer types except LAML and SKCM, only primary tumor specimens (TCGA code: 01) were analyzed. For the analysis of SKCM data, both primary and metastatic samples (TCGA code: 06) were analyzed. In the event that both a primary specimen and a metastatic specimen were available for SKCM, only the primary specimen was analyzed. For the analysis of LAML data, blood cancer specimens were given the TCGA code “03” and cancers with this code were included.

TCGA patient survival data and final clinical annotations were acquired from ref^40^. Selection of the clinical endpoint for each cancer type was based on the recommendations provided by ref^40^ based on data quality, cohort size, and the number of events that were observed. For the analysis of XIST expression, RPS4Y1 expression and patient sex, the TCGT cohort was excluded due to the reactivation of XIST that has been previously reported in testicular cancers^94^.

Pan-cancer oncogenes, oncogene-cancer type pairs, and recurrently-observed point mutations were acquired from ref^47^. Tumor mitotic activity and prostate cancer Gleason scores were acquired from the provisional TCGA annotations available at www.cbioportal.org95.

The list of FDA-approved cancer therapies was acquired from ref^72^. Drug targets were identified from the NCI drug dictionary^96^, from ref^97^, and from ref^98^. Multi-targeted kinase inhibitors (sorafenib, sunitinib, etc.) and other drugs where the mechanism-of-action is unclear were excluded from this analysis^99–101^. FDA approval dates were acquired from ref^102^ and ref^103^. Cancer dependency scores were acquired from www.depmap.org89.

Doubling times for the NCI-60 cell line panel were acquired from ref^104^. Suz12 binding sites in embryonic stem cells were acquired from ref^61^.

### Survival analysis in TCGA cohorts

The TCGA project was initiated to facilitate the molecular characterization of the major cancer types found in the US. While clinical and pathological data were collected for each patient, genomic analysis was prioritized over clinical follow-up. As justified by the analyses described in this manuscript and in other publications, we posit that performing survival analysis on the TCGA cohorts remains appropriate for several reasons. First, as described in Liu et al., the TCGA clinical data has been rigorously reviewed, harmonized, and validated through independent analyses^40^. Liu et al. also reported that, as expected, stage III/IV tumors in TCGA had uniformly worse outcomes compared to stage I/II tumors, and the median survival times for certain cancers fall within established ranges based on published case series. Secondly, in this work, we demonstrate that high-grade tumors have worse outcomes than low-grade tumors, that older patients have worse outcomes than younger patients, and that the survival times within TCGA cancer types are highly-correlated with the survival times reported in the nationally-representative SEER database (Figure S1). Thirdly, our analysis has verified the prognostic power of several established molecular biomarkers, including the adverse effects of p53 mutations and the strong association between cell cycle gene expression and aggressive disease^25, 27, 28, 54, 105^. Finally, we and others have verified that the frequencies of mutations in specific oncogenes and tumor suppressors in TCGA are very close to the frequencies observed in other clinical series, further demonstrating that the TCGA cohorts are broadly representative of cancer patients as a whole^29, 43, 106, 107^. Thus, while facilitating prognostic biomarker discovery was not the major goal of the TCGA project, we believe that the TCGA populations are representative cohorts and that the survival analysis we have conducted is appropriate.

### Selection of analysis methodology

Several statistical techniques have been developed to perform survival analysis^108^. In this paper, we chose to apply Cox proportional hazards regression to study the TCGA cohorts. The Cox model is given by the following function:

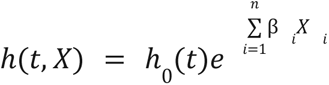

Where t is the survival time, h(t, X) is the hazard function, h_0_(t) is the baseline hazard, X_i_ is a potential prognostic variable, and β_i_ indicates the strength of the association between a prognostic variable and survival. In this model, patients have a baseline, time-dependent risk of death h_0_(t), modified by time-independent prognostic features that either increase (β_i_>0) or decrease (β_i_<0) risk of death. In this work, we report Z scores from these Cox models, which are calculated by dividing the regression coefficient (β_i_) by its standard error.

As we have previously described^29^, we utilize Cox proportional hazards modeling for survival analysis for several reasons. First, unlike Kaplan-Meier analysis, Cox models do not require the selection of threshold values, so continuous data like gene expression measurements do not need to be dichotomized. Secondly, Cox models can accept both continuous and discrete input data, allowing this approach to be used to analyze both binary (e.g., mutant vs. non-mutant) and continuous (e.g., gene expression) genomic features. Thirdly, Cox models can be used to perform both univariate (i = 1) and multivariate (i > 1) analyses. Fourthly, Cox regression allows us to calculate a Z score and a p value for each association, as Z scores represent the number of standard deviations from the mean of a normal distribution. Previous qq-analysis has demonstrated the underlying normality of the survival data^29^. Fifthly, Z scores encode the directionality of an association: poor prognostic factors will exhibit βi values greater than 0, while favorable prognostic factors will exhibit βi values less than 0. This allows “favorable” and “adverse” survival features to be directly compared. Sixthly, Z scores are useful for meta-analyses, as they can be combined using Stouffer’s Method^48^:

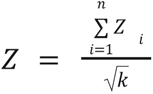

Seventhly, Cox proportional hazards modeling is commonly used in both previous genome-wide survival analyses and in numerous clinical biomarkers studies, facilitating comparison with other biomarker discovery efforts^21, 22, 29, 30, 109, 110^.

### Overall analysis strategy

In this paper, we describe the comprehensive and unbiased generation of Cox proportional hazard model Z scores for every genomic feature available in TCGA (CNAs, methylation, mutation, gene expression, miRNA expression, and protein expression), and for every cancer type. All analysis was performed in Python, using pandas^111^, matplotlib^112^, numpy^113^ and scipy^114^. Cox proportional hazards were computed using the R survival package^115^, and rpy2^116^ was used to integrate the R computations with Python scripts.

The software for these analyses is structured to be repeatable, modular, and debuggable. The same code was used for computing all Z scores, with swappable functions for preparing and cleaning each data type. Similarly, all clinical data was prepared identically across all analyses, using a custom-built parser for the clinical data provided by ref^40^.

TCGA copy number data was generated as relative copy number values for particular chromosomal intervals. This data was translated to produce a single copy number value on a per-gene basis, based on the observed copy number at each gene’s transcription start site. This annotation was performed using mapping data from GENCODE v32^117^. Interval trees, from the Python package intervaltree, were used to facilitate efficient mapping of segmental data to the appropriate genes^118^. The copy number value at a gene’s transcription start site was used as the input for the Cox models. Note that Cox proportional hazards modeling is threshold-independent, and so no minimum or maximum copy number value was required to specify a deletion or an amplification.

For protein expression data, the normalized and batch-corrected RPPA expression values were used as inputs for the Cox models. For the gene expression and microRNA expression data, values were log2-transformed and clipped at 0, then used as inputs for the Cox models.

For the methylation data, each probe was mapped to the relevant gene(s) that it recognized using the probeset annotations provided by Illumina. Beta values that mapped to the same gene were collapsed by averaging. This single average Beta value was used as input for the Cox models.

For CNAs, methylation, gene expression, miRNA expression, and protein expression, Cox models were only generated if at least 10 patients in a cohort had data for a particular feature.

For mutations, Cox models were generated if 2% or more of all sequenced patients for a particular cancer type had a non-synonymous mutation in the relevant gene. This 2% cut-off is based on qq tests for normality that we have previously conducted^29^. Non-synonymous mutations included: missense, nonsense, frameshift deletion, splice site, frameshift insertion, in-frame deletion, translation start site, nonstop mutation, and in-frame insertion. For each gene in each patient, a gene was considered to be mutated if there was a single of these non-synonymous mutations at any codon within the gene. In the driver gene analysis, a patient was marked as mutated if there was a single non-synonymous mutation at any of the relevant codons.

Multivariate analysis was performed using age, sex, stage, and grade data from ref.^40^ The variables used for each cohort are listed in Table S2G. The divisions for “stage” as a variable are listed in Table S2H. The divisions for “grade” as a variable are listed in Table S2I.

Single apostrophes were prepended to gene names in the output files from these analyses in order to allow the data tables to be read in Microsoft Excel without auto-formatting^119^.

### Kaplan-Meier analysis

Kaplan-Meier plots were generated using GraphPad Prism. Gene expression, miRNA expression, protein expression, and methylation values were dichotomized based on their mean values within the indicated cohorts. For copy number analysis, CNA values >0.3 were classified as amplified and CNA values <-0.3 were classified as deleted^29^.

### Gene ontology analysis

Gene ontology and transcription factor enrichment analysis were performed using g:Profiler^120^. Genes used to calculate a cell cycle gene score and transcription factor methylation score were also identified using the appropriate GO term via g:Profiler.

### Additional tools and resources

Gene set permutations were performed in Python by sampling 1000 random permutations of column data, in which observed Z scores were randomly assigned to gene or feature labels within each cancer type.

Peak finding was performed using the standard scipy signals library^114^. Gene network analysis for mitotic genes and developmental transcription factors was performed using NetworkX and _STRING_^121,122^.

### Code availability

The code used to perform the analysis in this paper is available at github.com/joan-smith/comprehensive-tcga-survival.

## Supporting information

Table S1. Univariate Cox Models

Table S2. Multivariate Cox Models

Table S3. Cox models with p53 status

Table S4. GO term analysis

Table S5. CNA peaks and valleys

Table S6. Univariate Cox models with recurrent mutations

Table S7. Drug targets

## Acknowledgments

Research in the Sheltzer Lab is supported by an NIH Early Independence award (1DP5OD021385), NIH grant R01CA237652, Department of Defense grant W81XWH-20-1-068, a Damon Runyon-Rachleff Innovation award, an American Cancer Society Research Scholar Grant, and a grant from the New York Community Trust.

## Author Contributions

J.C.S. and J.M.S. conceived, designed, and performed the analysis described in this work.

J.C.S. and J.M.S. wrote the manuscript and prepared the figures.

## Supplemental Tables

**Table S1. Genome-wide univariate Cox proportional hazard models to assess the link between individual genetic features and cancer patient outcomes.**

**Table S2. Genome-wide multivariate Cox proportional hazard models incorporating age, sex, stage, and grade to assess the link between individual genetic features and cancer patient outcomes.**

**Table S3. Genome-wide multivariate Cox proportional hazard models incorporating TP53 mutation status to assess the link between individual genetic features and cancer patient outcomes.**

**Table S4. Gene ontology enrichment among genetic features associated with cancer patient outcomes.**

**Table S5. Peaks and valleys identified in a topological analysis of copy number alteration Z scores by chromosomal coordinate.**

**Table S6. Univariate Cox proportional hazard models to assess the link between recurrent driver mutations and cancer patient outcomes.**

**Table S7. Analysis of the prognostic correlates of the targets of FDA-approved cancer drugs.**

---

**Figure S1.**
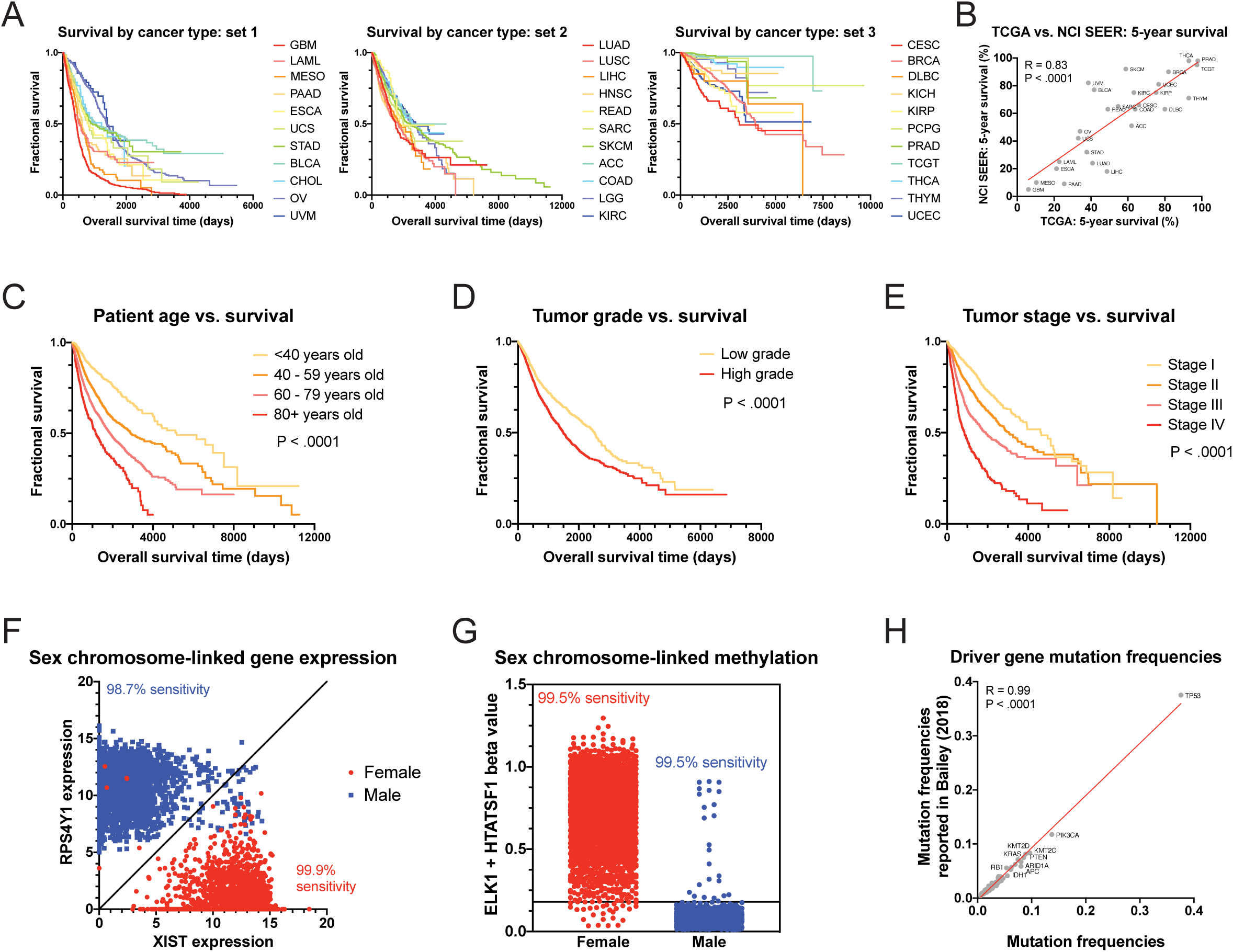
Validation of the genomic and clinical analysis pipeline. (A) Kaplan-Meier plots showing patient survival in the indicated TCGA cohorts. Patient cohorts were split into three groups based on average survival time. (B) A scatter plot comparing the 5-year survival frequency for the indicated TCGA cohorts with the 5-year survival frequency from the Surveillance, Epidemiology, and End Results (SEER) progra (C) Kaplan-Meier plot showing patient survival across all cancer types split according to patient age. (D) Kaplan-Meier plot showing patient survival across all cancer types split according to tumor grade. (E) Kaplan-Meier plot showing patient survival across all cancer types split according to tumor stage. (F) Scatter plot showing the expression of the X chromosome marker XIST transcript and the Y chromosome marker RPS4Y1. Nearly all female patients have high XIST expression and low RPS4Y1 expression, while nearly all male patients have high RPS4Y1 expression and low XIST expression, validating the link between gene expression profiles and clinical annotations. (G) Plot showing the methylation of the X chromosome genes ELK1 and HTATSF1. Nearly all female patients have high methylation levels of both genes, while nearly all male patients have low methylation levels of both genes, validating the link between methylation profiles and clinical annotations. (H) Scatter plot comparing the mutation frequency across cancer types of verified oncogenes calculated according to our analysis (X axis) or as reported in Bailey et al.^47^ (Y axis). The red line represents a linear regression plotted against the data.

**Figure S2.**
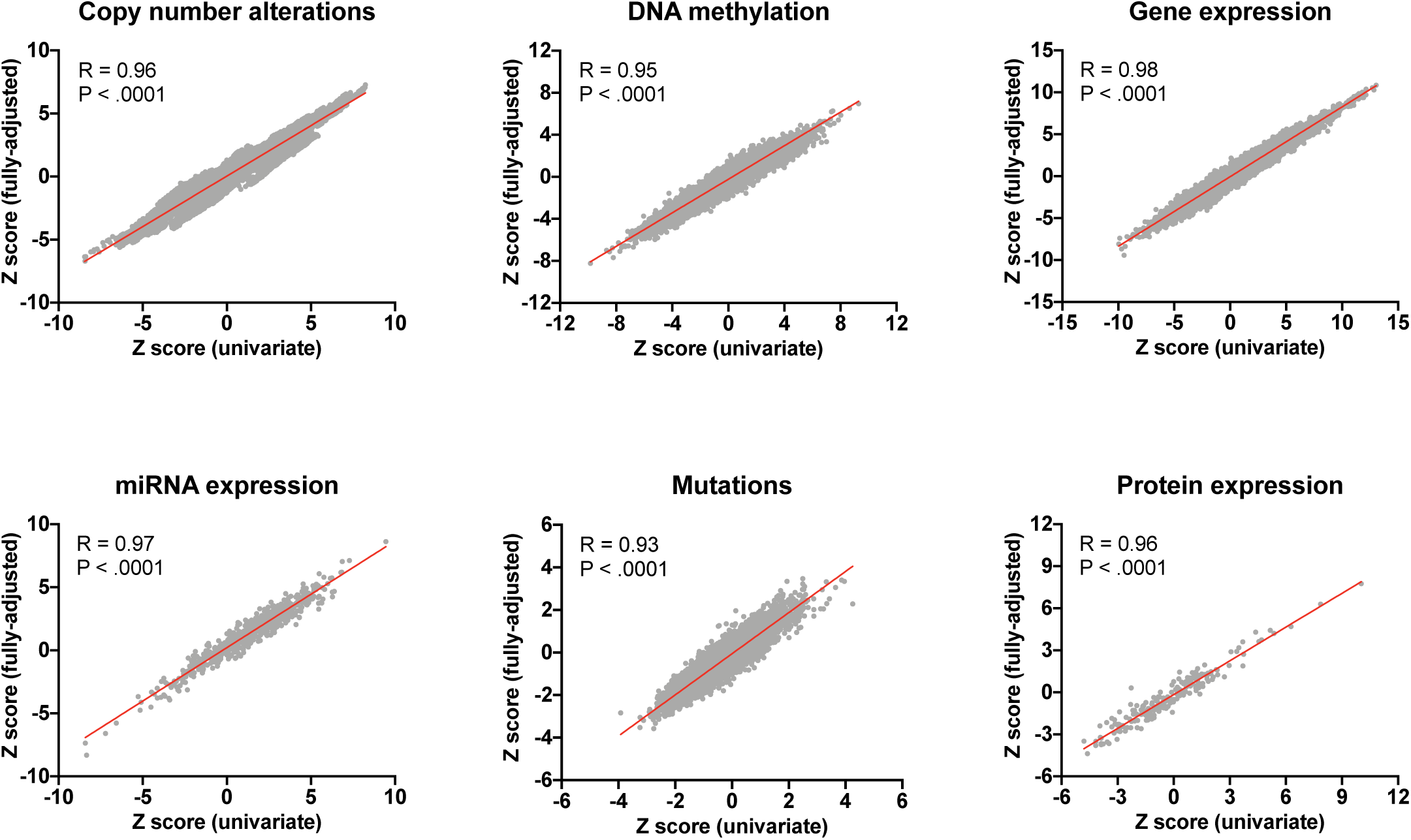
Univariate and multivariate (fully-adjusted) Cox proportional hazards models produce similar Z scores. Scatter plots comparing Stouffer’s Z from univariate models (X axis) and multivariate models (Y axis) for each of the six genomic platforms. The red lines represent linear regressions plotted against the data.

**Figure S3.**
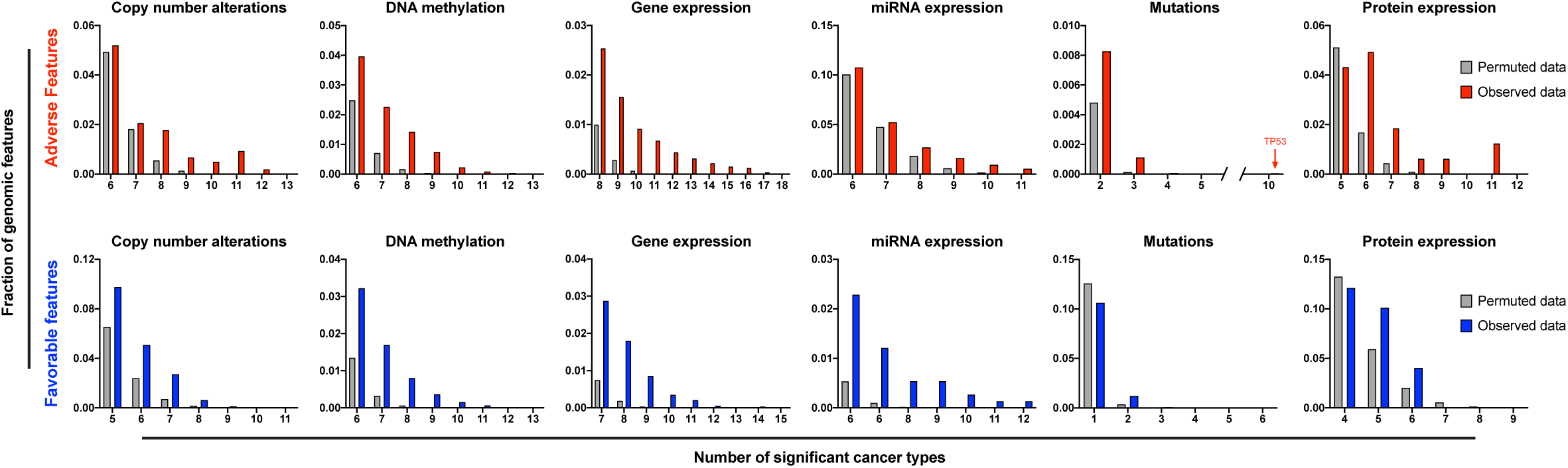
Some genomic features display significant correlations with patient outcome across cancer types. Histograms comparing the number of shared prognostic biomarkers across cancer types with values calculated by randomly permuting gene identity are displayed.

**Figure S4.**
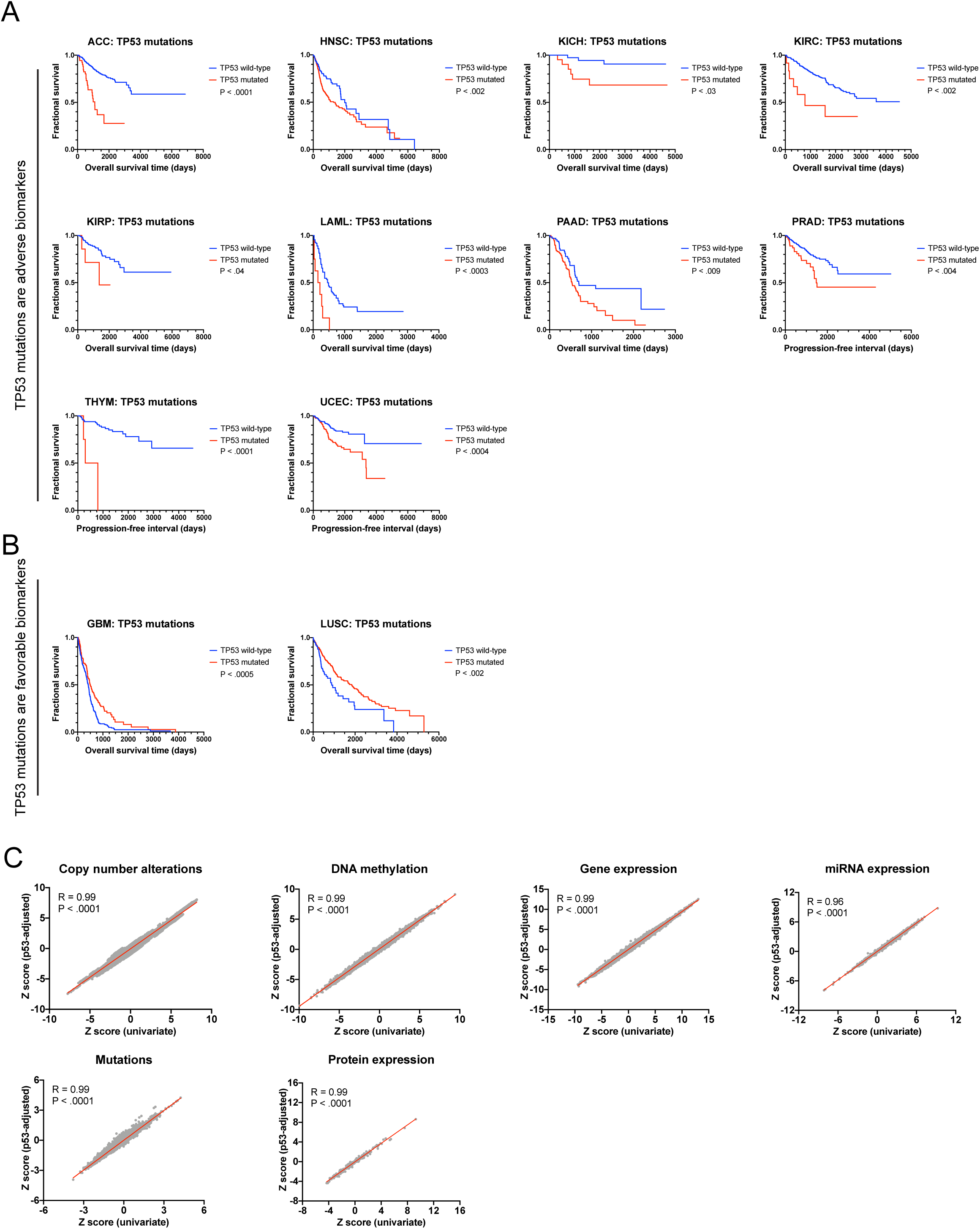
The prognostic significance of mutations in TP53. (A) Kaplan-Meier plots displaying the 10 cancer types in which mutations in TP53 are significantly associated with shorter survival times. (B) Kaplan-Meier plots displaying the two cancer types in which mutations in TP53 are significantly associated with longer survival times. (C) Scatter plots comparing Stouffer’s Z from univariate models (X axis) and multivariate models incorporating p53 status (Y axis) for each of the six genomic platforms. The red lines represent linear regressions plotted against the data.

**Figure S5.**
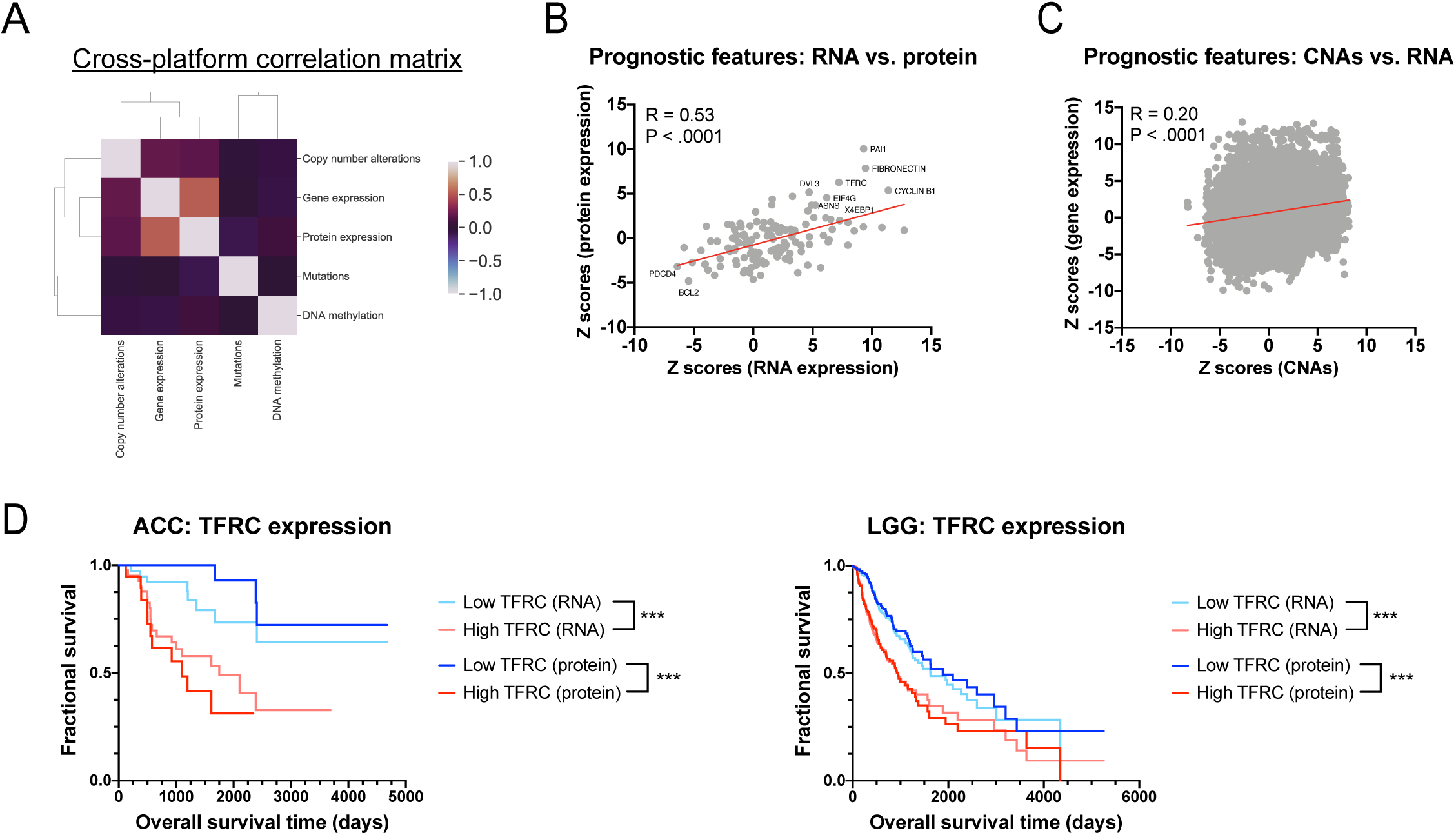
Correlation of prognostic features across genomic platforms. (A) A correlation matrix displaying the Pearson Correlation Coefficient between Z scores measured for the same gene on each platform. While gene expression Z scores and protein expression Z scores are strongly correlated (R = 0.53), the correlation between every other combination of platforms is significantly lower. (B) A scatter plot displaying Z scores calculated for gene expression (X axis) and Z scores calculated based on the expression of the same proteins (Y axis). The red line represents a linear regression plotted against the data. (C) A scatter plot displaying Z scores calculated for gene copy number (X axis) and Z scores calculated based on the expression of the same gene (Y axis). The red line represents a linear regression plotted against the data. (D) Kaplan-Meier plots displaying patient survival in ACC (left) and LGG (right) based on the expression of TFRC at either the RNA level or the protein level.

**Figure S6.**
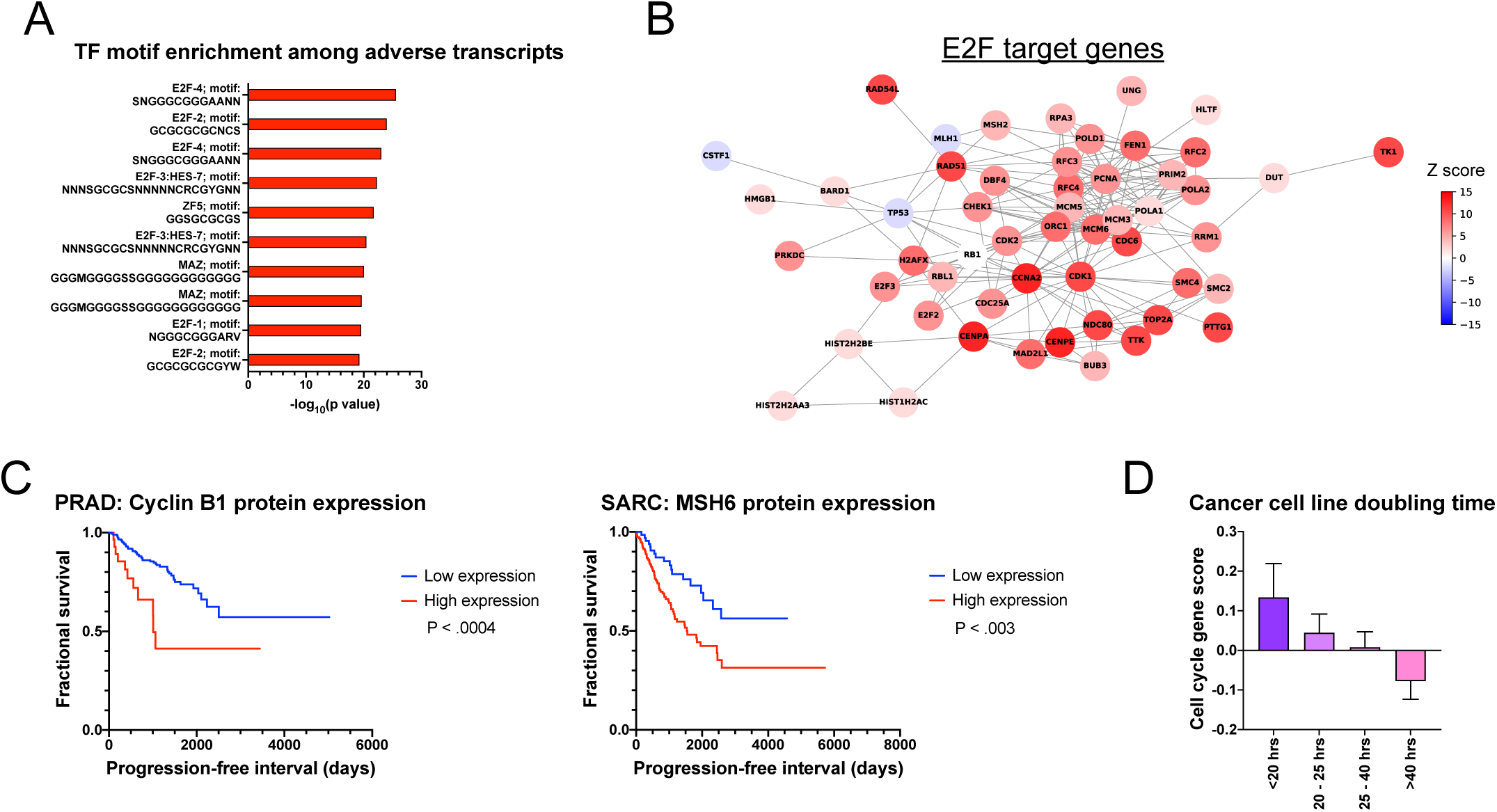
Transcripts and proteins associated with cell cycle progression are adverse prognostic features. (A) Enrichment of cell cycle-associated transcription factors among genes identified as adverse prognostic features. (B) Network of interactions among E2F target genes, colored according to Stouffer’s Z from the combined Cox models. (C) Kaplan-Meier plots showing that overexpression of the cell cycle-associated proteins Cyclin B1 (left) and MSH6 (right) are correlated with shorter survival times in PRAD and SARC, respectively. (D) Graph showing the correlation between cell cycle gene scores and the measured doubling times of the NCI-60 cancer cell line panel.

**Figure S7.**
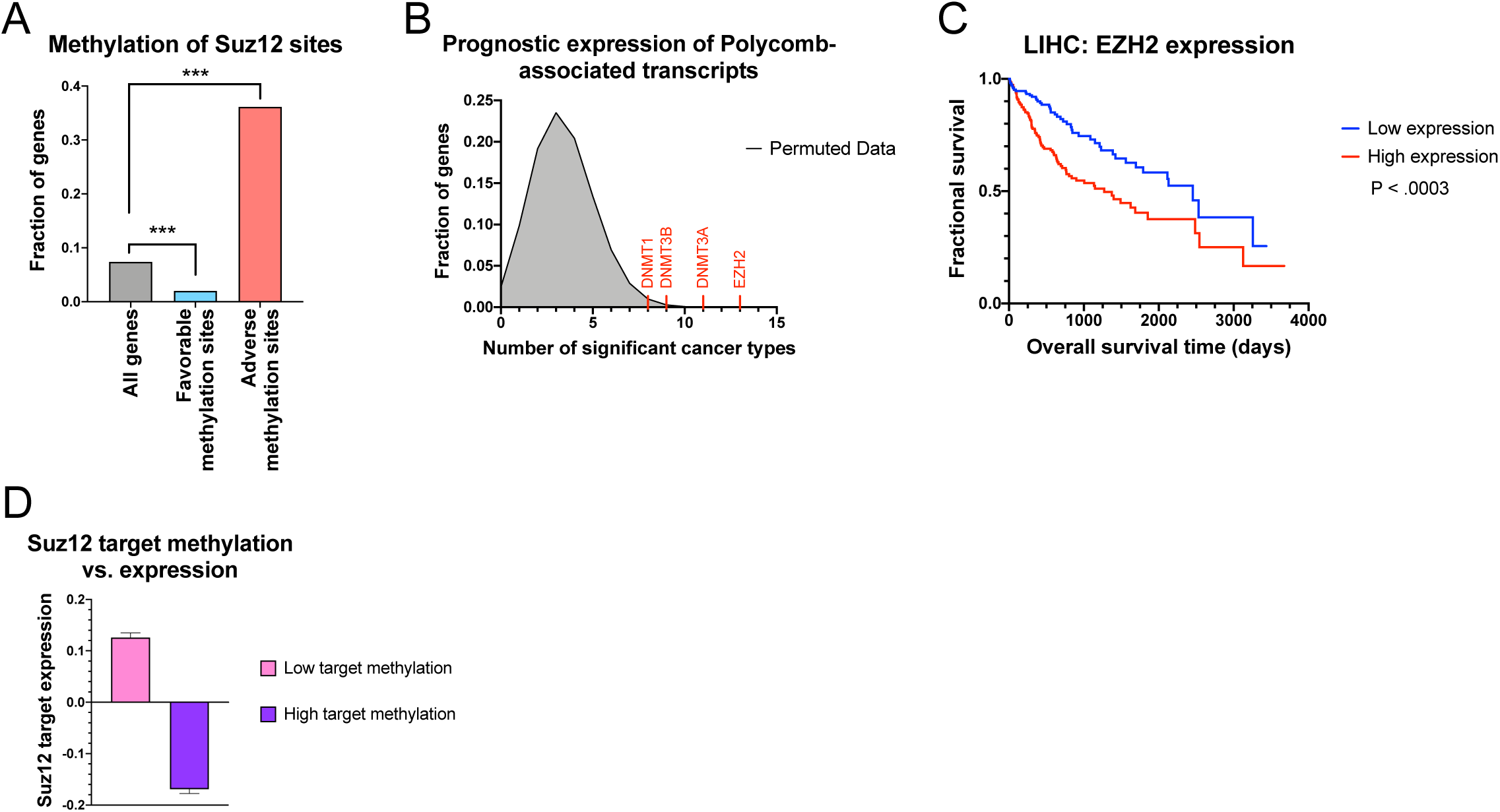
Overexpression of Polycomb components and methylation of Polycomb targets are pan-cancer adverse prognostic markers. (A) Adverse methylation sites are significantly enriched for genes bound by Polycomb-component Suz12 in hES cells. ***, P < .0005 (hypergeometric test). (B) Density plot showing the location of Polycomb components among genes identified as adverse features across multiple cancer types. (C) Kaplan-Meier plot showing that overexpression of the Polycomb component EZH2 is associated with shorter survival times in LIHC. (D) Graph showing the correlation between methylation levels at Polycomb target genes and decreased expression at those genes.

**Figure S8.**
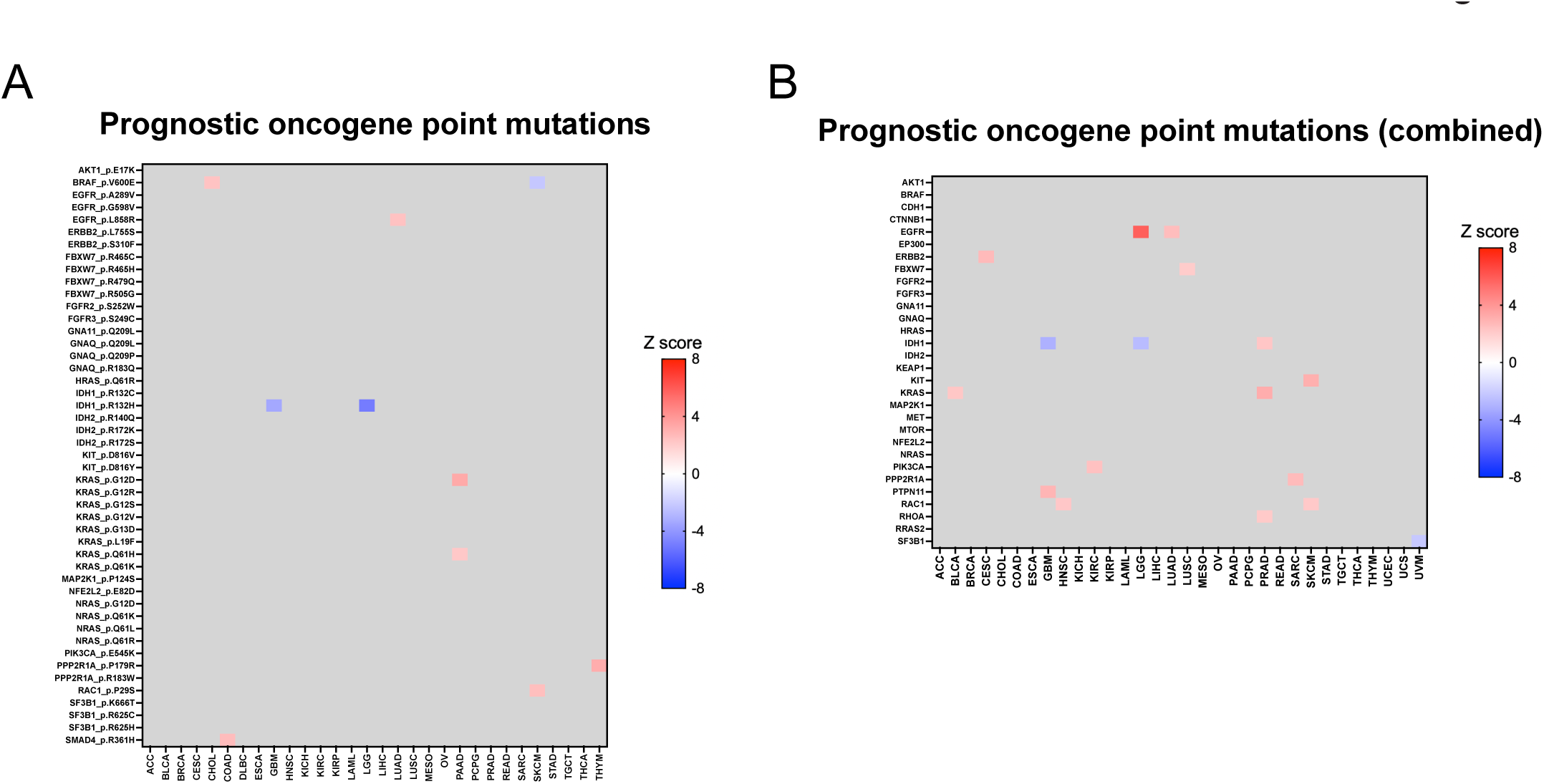
Recurrent point mutations are not widely associated with cancer patient outcome. (A) A heatmap showing significant (|Z| > 1.96) survival associations for recurrently-observed point mutations in oncogenes. Each row represents a specific mutation and each column represents a cancer patient cohort from TCGA. (B) A heatmap showing significant (|Z| > 1.96) survival associations for combinations of recurrently-observed point mutations in oncogenes. Each row represents a specific oncogene and each column represents a cancer patient cohort from TCGA.

**Figure S9.**
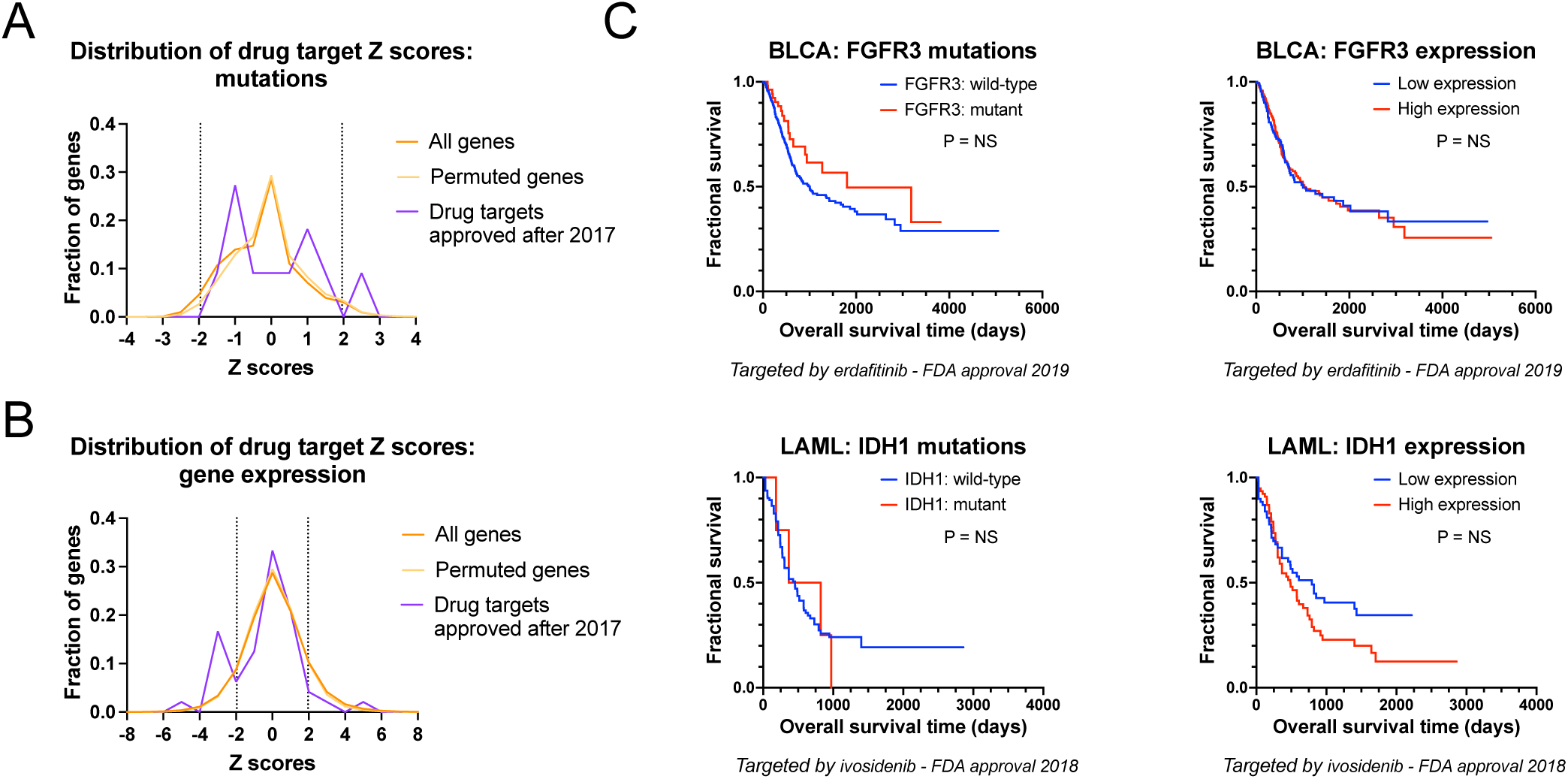
The targets of drugs that received FDA approval in 2018 or later are not strong prognostic biomarkers. (A) A density plot showing the distribution of mutation Z scores for the indicated gene sets. The dotted line at Z = −1.96 corresponds to P < .05 for a favorable mutation, while the dotted line at Z = 1.96 corresponds to P < .05 for an adverse mutation. (B) A density plot showing the distribution of gene expression Z scores for the indicated gene sets. The dotted line at Z = −1.96 corresponds to P < .05 for favorable overexpression, while the dotted line at Z = 1.96 corresponds to P < .05 for adverse overexpression. (C) Kaplan-Meier plots displaying survival times in BLCA (top) and LAML (bottom) based on the expression or mutation of FGFR3 (top) or IDH1 (bottom). The FGFR3 inhibitor erdafitinib received FDA approval for use in bladder cancer in 2019, while the IDH1 inhibitor ivosidenib received FDA approval for use in leukemia in 2018.

## Works Cited

1. Ludwig, J. A. & Weinstein, J. N. Biomarkers in Cancer Staging, Prognosis and Treatment Selection. Nat. Rev. Cancer 5, 845–856 (2005).

2. Dale, D. C. Poor prognosis in elderly patients with cancer: the role of bias and undertreatment. J. Support. Oncol. 1, 11–17 (2003).

3. Bouchardy, C. et al. Undertreatment Strongly Decreases Prognosis of Breast Cancer in Elderly Women. J. Clin. Oncol. 21, 3580–3587 (2003).

4. Bouchardy, C., Rapiti, E., Blagojevic, S., Vlastos, A.-T. & Vlastos, G. Older Female Cancer Patients: Importance, Causes, and Consequences of Undertreatment. J. Clin. Oncol. 25, 1858–1869 (2007).

5. Esserman, L. J., Thompson, I. M. & Reid, B. Overdiagnosis and overtreatment in cancer: an opportunity for improvement. JAMA 310, 797–798 (2013).

6. Jegerlehner, S. et al. Overdiagnosis and overtreatment of thyroid cancer: A population-based temporal trend study. PLOS ONE 12, e0179387 (2017).

7. Loeb, S. et al. Overdiagnosis and Overtreatment of Prostate Cancer. Eur. Urol. 65, 1046–1055 (2014).

8. Connolly, J. L., et al. Principles of Cancer Pathology. (2003).

9. Lang, H. et al. Multicenter determination of optimal interobserver agreement using the Fuhrman grading system for renal cell carcinoma: Assessment of 241 patients with > 15-year follow-up. Cancer 103, 625–629 (2005).

10. Griffiths, D. F. R. et al. A study of Gleason score interpretation in different groups of UK pathologists; techniques for improving reproducibility. Histopathology 48, 655–662 (2006).

11. Ozkan, T. A. et al. Interobserver variability in Gleason histological grading of prostate cancer. Scand. J. Urol. 50, 420–424 (2016).

12. Evans, A. J. et al. Interobserver Variability Between Expert Urologic Pathologists for Extraprostatic Extension and Surgical Margin Status in Radical Prostatectomy Specimens. Am. J. Surg. Pathol. 32, 1503–1512 (2008).

13. Gilks, C. B., Oliva, E. & Soslow, R. A. Poor interobserver reproducibility in the diagnosis of high-grade endometrial carcinoma. Am. J. Surg. Pathol. 37, 874–881 (2013).

14. Elmore, J. G. et al. Diagnostic Concordance Among Pathologists Interpreting Breast Biopsy Specimens. JAMA 313, 1122–1132 (2015).

15. Zaniboni, A. & Labianca, R. Adjuvant therapy for stage II colon cancer: an elephant in the living room? Ann. Oncol. 15, 1310–1318 (2004).

16. Bijker, N., Donker, M., Wesseling, J., Heeten, G. J. den & Rutgers, E. J. T. Is DCIS Breast Cancer, and How Do I Treat it? Curr. Treat. Options Oncol. 14, 75–87 (2013).

17. Young, R. C. Early-Stage Ovarian Cancer: To Treat or Not To Treat. JNCI J. Natl. Cancer Inst. 95, 94–95 (2003).

18. Berger, M. F. & Mardis, E. R. The emerging clinical relevance of genomics in cancer medicine. Nat. Rev. Clin. Oncol. 15, 353–365 (2018).

19. Sparano, J. A. et al. Adjuvant Chemotherapy Guided by a 21-Gene Expression Assay in Breast Cancer. N. Engl. J. Med. 379, 111–121 (2018).

20. Goossens, N., Nakagawa, S., Sun, X. & Hoshida, Y. Cancer biomarker discovery and validation. Transl. Cancer Res. 4, 256–269 (2015).

21. Gentles, A. J. et al. The prognostic landscape of genes and infiltrating immune cells across human cancers. Nat. Med. 21, 938–945 (2015).

22. Anaya, J., Reon, B., Chen, W.-M., Bekiranov, S. & Dutta, A. A pan-cancer analysis of prognostic genes. PeerJ 3, e1499 (2016).

23. Uhlen, M. et al. A pathology atlas of the human cancer transcriptome. Science 357, eaan2507 (2017).

24. Tang, Z., Kang, B., Li, C., Chen, T. & Zhang, Z. GEPIA2: an enhanced web server for large-scale expression profiling and interactive analysis. Nucleic Acids Res. 47, W556–W560 (2019).

25. Cuzick, J. et al. Prognostic value of an RNA expression signature derived from cell cycle proliferation genes in patients with prostate cancer: a retrospective study. Lancet Oncol. 12, 245–255 (2011).

26. Dancik, G. M. & Theodorescu, D. The Prognostic Value of Cell Cycle Gene Expression Signatures in Muscle Invasive, High-Grade Bladder Cancer. Bladder Cancer 1, 45–63 (2015).

27. Mosley, J. D. & Keri, R. A. Cell cycle correlated genes dictate the prognostic power of breast cancer gene lists. BMC Med. Genomics 1, 11 (2008).

28. Venet, D., Dumont, J. E. & Detours, V. Most Random Gene Expression Signatures Are Significantly Associated with Breast Cancer Outcome. PLOS Comput. Biol. 7, e1002240 (2011).

29. Smith, J. C. & Sheltzer, J. M. Systematic identification of mutations and copy number alterations associated with cancer patient prognosis. eLife 7, e39217 (2018).

30. Hieronymus, H. et al. Tumor copy number alteration burden is a pan-cancer prognostic factor associated with recurrence and death. eLife 7, (2018).

31. Stopsack, K. H. et al. Aneuploidy drives lethal progression in prostate cancer. Proc. Natl. Acad. Sci. U. S. A. 116, 11390–11395 (2019).

32. Vasudevan, A. et al. Single-Chromosomal Gains Can Function as Metastasis Suppressors and Promoters in Colon Cancer. Dev. Cell 52, 413–428.e6 (2020).

33. Vasudevan, A. et al. Aneuploidy as a promoter and suppressor of malignant growth. Nat. Rev. Cancer 21, 89–103 (2021).

34. Rosenthal, R. The file drawer problem and tolerance for null results. Psychol. Bull. 86, 638–641 (1979).

35. Shields, P. G. Publication Bias Is a Scientific Problem with Adverse Ethical Outcomes: The Case for a Section for Null Results. Cancer Epidemiol. Prev. Biomark. 9, 771–772 (2000).

36. Andre, F. et al. Biomarker studies: a call for a comprehensive biomarker study registry. Nat. Rev. Clin. Oncol. 8, 171–176 (2011).

37. The Cancer Genome Atlas Research Network et al. The Cancer Genome Atlas Pan-Cancer analysis project. Nat. Genet. 45, 1113–1120 (2013).

38. Anaya, J. OncoRank: A pan-cancer method of combining survival correlations and its application to mRNAs, miRNAs, and lncRNAs. https://peerj.com/preprints/2574 (2016) doi:10.7287/peerj.preprints.2574v1.

39. Tang, Z. et al. GEPIA: a web server for cancer and normal gene expression profiling and interactive analyses. Nucleic Acids Res. 45, W98–W102 (2017).

40. Liu, J. et al. An Integrated TCGA Pan-Cancer Clinical Data Resource to Drive High-Quality Survival Outcome Analytics. Cell 173, 400–416.e11 (2018).

41. Kaelin, W. G. Common pitfalls in preclinical cancer target validation. Nat. Rev. Cancer 17, 425–440 (2017).

42. Chopra, R. & Raynaud, F. I. Preclinical Studies to Enable First in Human Clinical Trials. in Phase I Oncology Drug Development (eds. Yap, T. A., Rodon, J. & Hong, D. S.) 45–69 (Springer International Publishing, 2020). doi:10.1007/978-3-030-47682-3_3.

43. Zehir, A. et al. Mutational landscape of metastatic cancer revealed from prospective clinical sequencing of 10,000 patients. Nat. Med. 23, 703–713 (2017).

44. Statistical Summaries - SEER Cancer Statistics. https://seer.cancer.gov/statistics/summaries.html.

45. Colonna, M., Bossard, N., Remontet, L. & Grosclaude, P. Changes in the risk of death from cancer up to five years after diagnosis in elderly patients: A study of five common cancers. Int. J. Cancer 127, 924–931 (2010).

46. AJCC Cancer Staging Manual. (Springer International Publishing, 2017).

47. Bailey, M. H. et al. Comprehensive Characterization of Cancer Driver Genes and Mutations. Cell 173, 371–385.e18 (2018).

48. Stouffer, S. A. The American soldier. (Princeton University Press, 1949).

49 Schmidt, >M. C. et al. Impact of genotype and morphology on the prognosis of glioblastoma. J. Neuropathol. Exp. Neurol. 61, 321–328 (2002).

50. Chen, Y.-J. et al. Association of Mutant TP53 with Alternative Lengthening of Telomeres and Favorable Prognosis in Glioma. Cancer Res. 66, 6473–6476 (2006).

51. Marks, J. R. et al. Overexpression and Mutation of p53 in Epithelial Ovarian Cancer. Cancer Res. 51, 2979–2984 (1991).

52. Rodrigues, N. R. et al. p53 mutations in colorectal cancer. Proc. Natl. Acad. Sci. 87, 7555–7559 (1990).

53. McShane, E. et al. Kinetic Analysis of Protein Stability Reveals Age-Dependent Degradation. Cell 167, 803–815.e21 (2016).

54. Baak, J. P. A. et al. Proliferation is the strongest prognosticator in node-negative breast cancer: significance, error sources, alternatives and comparison with molecular prognostic markers. Breast Cancer Res. Treat. 115, 241–254 (2009).

55. Whitfield, M. L., George, L. K., Grant, G. D. & Perou, C. M. Common markers of proliferation. Nat. Rev. Cancer 6, 99–106 (2006).

56. Black, A. R. & Azizkhan-Clifford, J. Regulation of E2F: a family of transcription factors involved in proliferation control. Gene 237, 281–302 (1999).

57. Sheltzer, J. M. A transcriptional and metabolic signature of primary aneuploidy is present in chromosomally-unstable cancer cells and informs clinical prognosis. Cancer Res. 73, 6401–6412 (2013).

58. Zhao, R., Choi, B. Y., Lee, M.-H., Bode, A. M. & Dong, Z. Implications of Genetic and Epigenetic Alterations of CDKN2A (p16(INK4a)) in Cancer. EBioMedicine 8, 30–39 (2016).

59. Schlesinger, Y. et al. Polycomb-mediated methylation on Lys27 of histone H3 pre-marks genes for de novo methylation in cancer. Nat. Genet. 39, 232–236 (2007).

60. Bracken, A. P. & Helin, K. Polycomb group proteins: navigators of lineage pathways led astray in cancer. Nat. Rev. Cancer 9, 773–784 (2009).

61. Lee, T. I. et al. Control of Developmental Regulators by Polycomb in Human Embryonic Stem Cells. Cell 125, 301–313 (2006).

62. Viré, E. et al. The Polycomb group protein EZH2 directly controls DNA methylation. Nature 439, 871–874 (2006).

63. Shinawi, T. et al. DNA methylation profiles of long- and short-term glioblastoma survivors. Epigenetics 8, 149–156 (2013).

64. Elias, A. D. Management of Small T1a/b N0 Breast Cancers. Am. Soc. Clin. Oncol. Educ. Book 10–19 (2012) doi:10.14694/EdBook_AM.2012.32.68.

65. Booth, C. M. et al. Adjuvant Chemotherapy for Stage II Colon Cancer: Practice Patterns and Effectiveness in the General Population. Clin. Oncol. 29, e29–e38 (2017).

66. Lee, K. et al. Adjuvant chemotherapy does not provide survival benefits to elderly patients with stage II colon cancer. Sci. Rep. 9, 11846 (2019).

67. Stark, J. R. et al. Gleason Score and Lethal Prostate Cancer: Does 3 + 4 = 4 + 3? J. Clin. Oncol. 27, 3459–3464 (2009).

68. Srigley, J. R. et al. Controversial issues in Gleason and International Society of Urological Pathology (ISUP) prostate cancer grading: proposed recommendations for international implementation. Pathology (Phila.) 51, 463–473 (2019).

69. Frederick, L., Wang, X.-Y., Eley, G. & James, C. D. Diversity and Frequency of Epidermal Growth Factor Receptor Mutations in Human Glioblastomas. Cancer Res. 60, 1383–1387 (2000).

70. Muñoz-Maldonado, C., Zimmer, Y. & Medová, M. A Comparative Analysis of Individual RAS Mutations in Cancer Biology. Front. Oncol. 9, (2019).

71. Hodis, E. et al. A Landscape of Driver Mutations in Melanoma. Cell 150, 251–263 (2012).

72. Cancer Drugs - National Cancer Institute. https://www.cancer.gov/about-cancer/treatment/drugs (2021).

73. Laszlo, G. S., Estey, E. H. & Walter, R. B. The past and future of CD33 as therapeutic target in acute myeloid leukemia. Blood Rev. 28, 143–153 (2014).

74. Gainor, J. F. et al. ALK rearrangements are mutually exclusive with mutations in EGFR or KRAS: an analysis of 1,683 patients with non-small cell lung cancer. Clin. Cancer Res. Off. J. Am. Assoc. Cancer Res. 19, 4273–4281 (2013).

75. Unni, A. M., Lockwood, W. W., Zejnullahu, K., Lee-Lin, S.-Q. & Varmus, H. Evidence that synthetic lethality underlies the mutual exclusivity of oncogenic KRAS and EGFR mutations in lung adenocarcinoma. eLife 4, e06907 (2015).

76. Mack, P. C. et al. Spectrum of driver mutations and clinical impact of circulating tumor DNA analysis in non–small cell lung cancer: Analysis of over 8000 cases. Cancer 126, 3219–3228 (2020).

77. Darvin, P., Toor, S. M., Sasidharan Nair, V. & Elkord, E. Immune checkpoint inhibitors: recent progress and potential biomarkers. Exp. Mol. Med. 50, 1–11 (2018).

78. Sun, L. et al. Clinical efficacy and safety of anti-PD-1/PD-L1 inhibitors for the treatment of advanced or metastatic cancer: a systematic review and meta-analysis. Sci. Rep. 10, 2083 (2020).

79. Singh, S. et al. The emerging role of immune checkpoint inhibitors in the treatment of triple-negative breast cancer. Drug Discov. Today (2021) doi:10.1016/j.drudis.2021.03.011.

80. Francisco, L. M., Sage, P. T. & Sharpe, A. H. The PD-1 pathway in tolerance and autoimmunity. Immunol. Rev. 236, 219–242 (2010).

81. Ahmadzadeh, M. et al. Tumor antigen–specific CD8 T cells infiltrating the tumor express high levels of PD-1 and are functionally impaired. Blood 114, 1537–1544 (2009).

82. Ali, H. R. et al. Association between CD8+ T-cell infiltration and breast cancer survival in 12 439 patients. Ann. Oncol. 25, 1536–1543 (2014).

83. Jarmula, A. Antifolate Inhibitors of Thymidylate Synthase as Anticancer Drugs. Mini Rev. Med. Chem. 10, 1211–1222 (2010).

84. Peters, G. J. et al. Induction of thymidylate synthase as a 5-fluorouracil resistance mechanism. Biochim. Biophys. Acta BBA - Mol. Basis Dis. 1587, 194–205 (2002).

85. Markham, A. Erdafitinib: First Global Approval. Drugs 79, 1017–1021 (2019).

86. Gutteridge, R. E. A., Ndiaye, M. A., Liu, X. & Ahmad, N. Plk1 inhibitors in cancer therapy: From laboratory to clinics. Mol. Cancer Ther. 15, 1427–1435 (2016).

87. Green, S. D. & Konig, H. Treatment of Acute Myeloid Leukemia in the Era of Genomics—Achievements and Persisting Challenges. Front. Genet. 11, (2020).

88. Aye, Y., Li, M., Long, M. J. C. & Weiss, R. S. Ribonucleotide reductase and cancer: biological mechanisms and targeted therapies. Oncogene 34, 2011–2021 (2015).

89. . Meyers, R. M. et al. Computational correction of copy number effect improves specificity of CRISPR-Cas9 essentiality screens in cancer cells. Nat. Genet. 49, 1779–1784 (2017).

90. DepMap, B. DepMap Achilles 18Q3 public. (2018) doi:10.6084/m9.figshare.6931364.v1.

91. Conway, J. R., Warner, J. L., Rubinstein, W. S. & Miller, R. S. Next-Generation Sequencing and the Clinical Oncology Workflow: Data Challenges, Proposed Solutions, and a Call to Action. JCO Precis. Oncol. 1–10 (2019) doi:10.1200/PO.19.00232.

92. William L Henrich and John M Burkat. Patient survival and maintenance dialysis. UpToDate (2021).

93. PanCanAtlas Publications | NCI Genomic Data Commons. https://gdc.cancer.gov/about-data/publications/pancanatlas.

94. Looijenga, L. H., Gillis, A. J., van Gurp, R. J., Verkerk, A. J. & Oosterhuis, J. W. X inactivation in human testicular tumors. XIST expression and androgen receptor methylation status. Am. J. Pathol. 151, 581–590 (1997).

95. Gao, J. et al. Integrative Analysis of Complex Cancer Genomics and Clinical Profiles Using the cBioPortal. Sci. Signal. 6, pl1 (2013).

96. NCI Drug Dictionary - National Cancer Institute. https://www.cancer.gov/publications/dictionaries/cancer-drug/ (2021).

97. Corsello, S. M. et al. Discovering the anticancer potential of non-oncology drugs by systematic viability profiling. *Nat*. Cancer 1, 235–248 (2020).

98. Wishart, D. S. et al. DrugBank 5.0: a major update to the DrugBank database for 2018. Nucleic Acids Res. 46, D1074–D1082 (2018).

99. Gentile, C., Martorana, A., Lauria, A. & Bonsignore, R. Kinase Inhibitors in Multitargeted Cancer Therapy. Curr. Med. Chem. 24, 1671–1686 (2017).

100. Lin, A. et al. Off-target toxicity is a common mechanism of action of cancer drugs undergoing clinical trials. Sci. Transl. Med. 11, eaaw8412 (2019).

101. Lin, A. & Sheltzer, J. M. Discovering and validating cancer genetic dependencies: approaches and pitfalls. Nat. Rev. Genet. 21, 671–682 (2020).

102. Hematology/Oncology (Cancer) Approvals & Safety Notifications. FDA https://www.fda.gov/drugs/resources-information-approved-drugs/hematologyoncology-cancer-approvals-safety-notifications (2021).

103. FDA Approval History. *Drugs.com* https://www.drugs.com/history/.

104. Reinhold, W. C. et al. CellMiner: A Web-Based Suite of Genomic and Pharmacologic Tools to Explore Transcript and Drug Patterns in the NCI-60 Cell Line Set. Cancer Res. 72, 3499–3511 (2012).

105. Robles, A. I. & Harris, C. C. Clinical Outcomes and Correlates of TP53 Mutations and Cancer. Cold Spring Harb. Perspect. Biol. 2, (2010).

106. Jordan, E. J. et al. Prospective Comprehensive Molecular Characterization of Lung Adenocarcinomas for Efficient Patient Matching to Approved and Emerging Therapies. Cancer Discov. 7, 596–609 (2017).

107. Amar, D., Izraeli, S. & Shamir, R. Utilizing somatic mutation data from numerous studies for cancer research: proof of concept and applications. Oncogene 36, 3375–3383 (2017).

108. Kleinbaum, D. G. & Klein, M. Survival Analysis: A Self-Learning Text, Third Edition. (Springer-Verlag, 2012).

109. Fukuoka, M. et al. Biomarker Analyses and Final Overall Survival Results From a Phase III, Randomized, Open-Label, First-Line Study of Gefitinib Versus Carboplatin/Paclitaxel in Clinically Selected Patients With Advanced Non–Small-Cell Lung Cancer in Asia (IPASS). J. Clin. Oncol. 29, 2866–2874 (2011).

110. Parker, J. S. et al. Supervised Risk Predictor of Breast Cancer Based on Intrinsic Subtypes. J. Clin. Oncol. 27, 1160–1167 (2009).

111. Jeff Reback et al. pandas-dev/pandas: Pandas 1.0.3. (Zenodo, 2020). doi:10.5281/zenodo.3715232.

112. Hunter, J. D. Matplotlib: A 2D Graphics Environment. Comput. Sci. Eng. 9, 90–95 (2007).

113. Harris, C. R. et al. Array programming with NumPy. Nature 585, 357–362 (2020).

114. Virtanen, P. et al. SciPy 1.0: fundamental algorithms for scientific computing in Python. Nat. Methods 17, 261–272 (2020).

115. Therneau, T. M. Survival Analysis [R package survival version 3.2-11]. https://CRAN.R-project.org/package=survival (2021).

116. rpy2/rpy2. https://github.com/rpy2/rpy2 (2021).

117. Frankish, A. et al. GENCODE reference annotation for the human and mouse genomes. Nucleic Acids Res. 47, D766–D773 (2019).

118. Halbert, C.-L. chaimleib/intervaltree. https://github.com/chaimleib/intervaltree (2021).

119. Zeeberg, B. R. et al. Mistaken Identifiers: Gene name errors can be introduced inadvertently when using Excel in bioinformatics. BMC Bioinformatics 5, 80 (2004).

120. Raudvere, U. et al. g:Profiler: a web server for functional enrichment analysis and conversions of gene lists (2019 update). Nucleic Acids Res. 47, W191–W198 (2019).

121. Proceedings of the Python in Science Conference (SciPy): Exploring Network Structure, Dynamics, and Function using NetworkX. http://conference.scipy.org/proceedings/SciPy2008/paper_2/.

122. Szklarczyk, D. et al. STRING v11: protein-protein association networks with increased coverage, supporting functional discovery in genome-wide experimental datasets. Nucleic Acids Res. 47, D607–D613 (2019).

